# Evolutionary adaptation to juvenile malnutrition impacts adult metabolism and impairs adult fitness in *Drosophila*

**DOI:** 10.1101/2022.01.11.475896

**Authors:** Berra Erkosar, Cindy Dupuis, Fanny Cavigliasso, Loriane Savary, Laurent Kremmer, Hector Gallart-Ayala, Julijana Ivanisevic, Tadeusz J. Kawecki

## Abstract

Juvenile undernutrition has lasting effects on adult metabolism of the affected individuals, but it is unclear how adult physiology is shaped over evolutionary time by natural selection driven by juvenile undernutrition. We combined RNAseq, targeted metabolomics and genomics to study the consequences of evolution under juvenile undernutrition for metabolism of reproductively active adult females of *Drosophila melanogaster*. Compared to Control populations maintained on standard diet, Selected populations maintained for over 230 generations on a nutrient-poor larval diet evolved major changes in adult gene expression and metabolite abundance, in particular affecting amino-acid and purine metabolism. The evolved differences in adult gene expression and metabolite abundance between Selected and Control populations were positively correlated with the corresponding differences previously reported for Selected versus Control larvae. This implies that genetic variants affect both stages similarly. Even when well fed, the metabolic profile of Selected flies resembled that of flies subject to starvation. Finally, Selected flies had lower reproductive output than Controls even when both were raised under the conditions under which the Selected populations evolved. These results imply that evolutionary adaptation to juvenile undernutrition has large pleiotropic consequences for adult metabolism, and that they are costly rather than adaptive for adult fitness. Thus, juvenile and adult metabolism do not appear to evolve independently from each other even in a holometabolous species where the two life stages are separated by a complete metamorphosis.

## Introduction

Nutrient shortage is an important and widespread ecological stressor, and a driver of natural selection on animal physiology. Juveniles are often the first to suffer from undernutrition, as they are usually competitively inferior to adults. Furthermore, in most species, juveniles have no option to arrest their development and wait for better times; they are forced to continue development despite famine. Prolonged nutrient shortage at a juvenile stage usually results in stunted adult size, but also often has profound effects on adult physiology that do not seem mediated by body size. They include changes in epigenetic marks (Lillycrop and Burdge 2015), gene expression (May and Zwaan 2017; Szostaczuk, et al. 2020) metabolome (Agnoux, et al. 2014), triglyceride accumulation (Lukaszewski, et al. 2013; Klepsatel, et al. 2020), blood pressure (de Brito Alves, et al. 2014), behavior (Akitake, et al. 2015) and longevity (Davis, et al. 2016). In humans, prenatal undernutrition is correlated with negative consequences for diverse aspects of metabolic function and health at old age (Roseboom, et al. 2006; de Rooij, et al. 2010).

These physiological responses to developmental nutritional conditions are a form of phenotypic plasticity; i.e., a change of phenotype induced by differences in the environment, with no change in genome sequence (Scheiner 1993; Bateson, et al. 2004). We know much less on whether and how adult physiology and metabolism evolve genetically over generations in response to natural selection driven by repeated exposure of the population to juvenile undernutrition. Some authors postulate that phenotypic plasticity in response to novel environmental conditions is usually adaptive, and that evolutionary change will mostly reinforce these initially plastic phenotypes (“genetic assimilation”) (Baldwin 1896; Pigliucci and Murren 2003). Thus, plasticity would predict evolution. However, this prediction does not appear borne out by empirical data. In particular, experimental studies that compared plastic and evolutionary responses of gene expression to novel environments usually found little overlap between genes involved in the two responses; for those that did overlap, evolutionary change tended to counter rather than be in line with plasticity (Yampolsky, et al. 2012; Ghalambor, et al. 2015; Huang and Agrawal 2016), for an exception see Josephs, et al. (2021). Thus, to understand how adult physiology is molded by exposure to juvenile undernutrition over evolutionary time, inferences based on phenotypically plastic responses should be complemented by the study of genetically-based adaptations.

The particular vulnerability of juveniles to famine also raises the question of the degree to which metabolism and physiology of different life stages can evolve independently if they face different nutritional conditions. Anurans and holometabolous insects demonstrate that in the long run evolution can generate spectacular differences in morphology and physiology between juvenile and adult stages. This has led some authors to argue that metamorphosis of holometabolous insects allows the larval and adult phenotypes to be evolutionarily decoupled from each other (Moran 1994; Rolff, et al. 2019). This postulate requires a sufficient supply of genetic variants that affect relevant phenotypes in a stage-specific manner. However, surprisingly little is known to what degree this is the case, whether in holometabolous insects or other animals (Collet and Fellous 2019). For example, while some enzymes in *Drosophila* have distinct larval and adult forms encoded by different genes, most metabolic genes are expressed at both stages (www.flybase.org). Thus, independent evolution of larval and adult metabolism would require independent regulation of expression of the same genes. It may be more parsimonious to expect that most regulatory genetic variants would tend to affect gene expression across lifetime, constraining independent evolution of the juvenile and adult stages. If so, response to selection driven by juvenile undernutrition would lead to similar physiological change in larvae and adults, and these changes might be disadvantageous for adult performance, in particular under conditions of plentiful food.

Experimental data to address the above questions are scarce. Experimental evolution studies on *Drosophila* demonstrated that adaptation to poor larval nutrition (nutrient-poor diet or larval crowding) may be associated with changes in adult traits such as fecundity (May, et al. 2019), lifespan (Shenoi, et al. 2016; May, et al. 2019), starvation resistance (Kawecki, et al. 2021), pathogen resistance (Vijendravarma, et al. 2015) or heat tolerance (Kapila, et al. 2021). Changes in gene expression and metabolism underlying these correlated responses of adult phenotypes to selection for juvenile or larval undernutrition tolerance have not been explored. Insights into if and how selection driven by juvenile undernutrition shapes adult metabolism would shed light on the importance of evolutionary forces acting on development for understanding adult physiology, with potential implications for human health (Neel 1962; Prentice, et al. 2005).

Here, we use experimental evolution to study the consequences of evolutionary adaptation to larval undernutrition for gene expression, metabolite abundance and reproductive performance of adult *Drosophila melanogaster* females. We compare six replicate populations (the “Selected” populations) evolved for over 230 generations on a very nutrient-poor larval diet, to six populations of the same origin maintained on a moderately nutrient-rich standard larval diet (the “Control” populations). Importantly, in the course of experimental evolution flies of both sets of populations were always transferred to fresh standard diet soon after emergence, maintained on this diet for 4-6 days, and additionally fed ad libitum live yeast before reproduction. Thus, every generation of Selected populations experienced a switch from the poor larval diet to the standard adult diet. Compared to the Controls, larvae of the Selected populations evolved higher viability, faster development and faster growth on the poor diet (Kolss, et al. 2009; Vijendravarma and Kawecki 2013; Cavigliasso, et al. 2020). The Selected larvae appear to invest more in protein digestion (Erkosar, et al. 2017), are more efficient in extracting amino acids from the poor diet (Cavigliasso, et al. 2020), and are able to initiate metamorphosis at a smaller size (Vijendravarma, et al. 2012), traits that likely contribute to their improved performance on the poor diet. The Selected populations also evolved reduced dependence on microbiota for development on poor diet (Erkosar, et al. 2017), However, they show greater susceptibility to an intestinal pathogen (Vijendravarma, et al. 2015) and are less resistant than Controls to starvation at the adult stage (Kawecki, et al. 2021). Finally, the Selected and Control larvae also show divergence in metabolite abundance consistent with evolutionary changes in amino acid metabolism, with lower free amino acid concentrations and a higher rate of amino acid catabolism (Cavigliasso, et al. 2023). Evolutionary divergence between Selected and Control populations is highly polygenic, involving over 100 genomic regions enriched in hormonal and metabolic genes (Kawecki, et al. 2021). Thus, evolutionary adaptation to larval undernutrition in our experimental populations involved a complex suite of genomic and phenotypic changes.

Motivated by the general questions discussed above, in this study we address the following specific questions about the evolutionary divergence between the Selected and Control populations at the adult stage:

First, have the Selected and Control populations evolved major genetically-based differences in adult gene expression and metabolite abundance, implying that evolutionary adaptation to larval undernutrition has significant consequences for the physiology of the adult stage? Which pathways have been affected?

Second, in view of the controversy about the relationship between plasticity and evolution summarized above, to what degree do these genetically-based evolutionary changes in adult gene expression and metabolome resemble the phenotypically plastic responses to poor larval diet?

Third, are differences in adult gene expression and metabolite abundance between Selected and Control populations similar to those between Selected and Control larvae (reported elsewhere; Erkosar, et al. 2017; Cavigliasso, et al. 2023)? Such a similarity would be expected either if both life stages were exposed to similar selection pressures, or if genetic variants favored by larval undernutrition had pleiotropic effects on gene expression at the two stages. As we argue in Discussion, we find the latter interpretation more parsimonious, even it contradicts the notion that larval and adult gene expression patterns can be decoupled from each other in holometabolous insect (Moran 1994; Rolff, et al. 2019).

Fourth, do the metabolic profiles of Selected and Control flies differ along a similar axis as the profiles of fed versus starving flies? Selected flies are less resistant to starvation than Controls and the genomic architecture of divergence between them shares many genes with starvation resistance (Kawecki, et al. 2021). This question addresses the link between these two findings, and the answer would aid in functional interpretation of metabolic changes evolved by the Selected populations.

Finally, do adult flies of the Selected populations perform better in terms of fitness than Controls when both are raised on the poor larval diet? This would be expected if most changes in their adult physiology evolved because they improve performance of adults raised on poor diet. In contrast, if the metabolic changes observed at the adult stage primarily reflect pleiotropic effects of genetic variants that improve larval growth and survival on the poor diet, Selected adults should not have an advantage or even be inferior to Controls even when raised under the conditions under which the former evolved.

## Results

### Gene expression patterns point to evolutionary changes in adult metabolism

To explore plastic and evolutionary responses of adult gene expression to poor diet we performed RNAseq on 4-day old adult females from the Selected and Control populations raised on either larval diet. In this factorial design, the effect of larval diet treatment (poor vs standard) on the mean expression level is a measure of the plastic response. The difference between Selected and Control populations assessed in flies raised on the same diet is a measure of genetically-based evolutionary change (Figure 1A). Potential differences of the plastic responses between Selected and Control populations or the influence of diet on the expression of evolutionary divergence would be reflected in the evolutionary regime × diet interaction.

**Figure 1.**
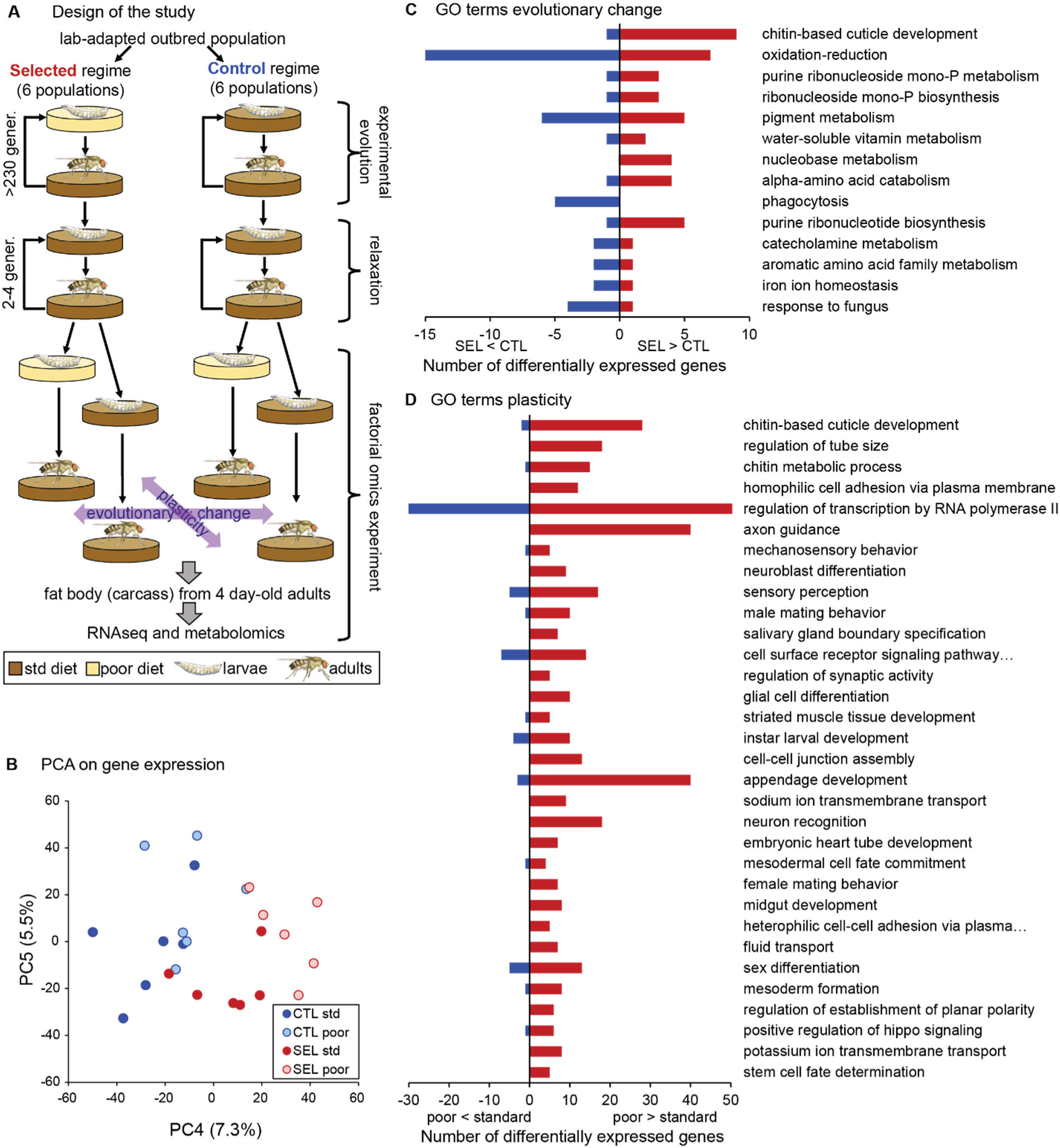
Phenotypic plasticity and genetically-based evolutionary change in response to larval undernutrition both lead to divergence in gene expression patterns of adult *Drosophila* females. (A) Design of the experimental evolution and of the gene expression assay. The purpose of “relaxation” was to eliminate potential effects of (grand)parental environment. The purple arrows indicate the two main factors in the analysis and interpretation, the evolutionary history (evolutionary change) and the effect of larval diet on which the current generation was raised (plasticity). (B) Sample score plot of the 4th and 5th principal components on the expression levels of 8701 genes of flies from Selected (SEL) and Control (CTL) populations raised on poor and standard (std) larval diet. Each point represents a replicate population × diet combination (Figure 1—source data 1). For complete plots of PC1-6 see Figure 1 - figure supplement 1. (C) Biological process GO terms enriched (at *P* < 0.01) in genes differentially expressed between Selected and Control populations, with the number of genes upregulated (red) and downregulated (blue) in Selected populations. (D) GO terms enriched at *P* < 0.01 among genes showing a plastic response to diet. For details of the GO-term enrichment results see Supplementary file 2.

Prior to collection for RNAseq, the flies were transferred to standard diet upon emergence from the pupa and maintained in mixed-sex group; thus, they were mated and reproductively active. We focused on female abdominal fat body, the key metabolic organ combining the functions of mammalian liver and adipose tissue. It is in the fat body where metabolic reserves of glycogen and triglycerides are stored and mobilized, and where dietary nutrients are converted into the proteins and lipids subsequently transported to the ovaries for egg production (Li, et al. 2019). Because a clean dissection of adult fat body is difficult, we performed RNAseq on the “carcass”, consisting of the fat body together with the attached abdominal body wall and any embedded neurons, but excluding the digestive, reproductive and most of the excretory organs.

Based on the estimation of true nulls (Storey and Tibshirani 2003), the evolutionary regime (i.e., Selected versus Control) affected adult expression of about 26 % of all 8701 genes included in the analysis while the larval diet affected about 45%. Allowing for 10% false discovery rate (FDR), we identified 219 genes as differentially expressed between Selected and Control populations and 827 between flies raised on standard or poor diet (Supplementary file 1). The magnitude of the changes in gene expression was moderate, with the significant genes showing on average about 1.76-fold difference (in either direction) between Selected and Control populations and about 1.30-fold difference between flies raised on poor versus standard diet. While the first three principal component axes appear driven by idiosyncratic variation among replicate populations and individual samples, PC4 and PC5 clearly differentiate the evolutionary regimes or larval diets (Figure 1B, Figure 1 – figure supplement 1). The divergence of gene expression patterns between Selected and Control populations, as well as between flies raised on poor and standard diet, was supported by MANOVA on the first five principal components, jointly accounting for 67% of variance (Wilks’ λ = 0.052, *F*_6,5_ = 15.1, *P* = 0.0046 for regime and Wilks’ λ = 0.057, *F*_6,5_ = 13.9, *P* = 0.0055 for diet). We detected no interaction between the evolutionary regime and current diet either in the PCA-MANOVA (Wilks’ λ = 0.59, *F*_6,5_ = 0.6, *P* = 0.41) or in the gene-by-gene analysis (lowest *q* = 99.9%). Thus, both the evolutionary adaptation and the phenotypically plastic response to larval diet experienced had substantial effects on adult gene expression patterns; however, these effects appeared largely additive (i.e., independent of each other).

The strongest statistical signal from gene ontology (GO) enrichment analysis on genes differentially expressed between Selected and Control flies points to genes involved in cuticle development and maturation, including several structural cuticle proteins (Figure 1C, Supplementary file 2 table A). Most of these genes are mainly expressed in epidermis (including in the tracheal system), and nearly all show higher expression in the Selected compared to Control populations. This may reflect differences in the relative amount of the epidermal tissues or in the rate of cuticle maturation. Several other enriched GO terms (phagocytosis, pigment metabolic process, iron ion homeostasis, response to fungus) include multiple genes expressed mainly in hemocytes; these genes are downregulated in Selected flies, suggesting lower abundance of hemocytes.

The other top enriched GO terms point to metabolic processes, notably oxidation/reduction, biosynthesis of purine compounds and amino acid catabolism (Figure 1C). Differentially expressed genes involved in amino acid catabolism included enzymes involved in catabolism of arginine (*Oat*), serine (*Shmt*), branched-chain amino acids (*CG17896, CG6638*, *Dbct*), tryptophan (*Trh*), asparagine (*CG7860*) and tyrosine (*CG1461*). Thus, the signal of enrichment in amino acid metabolism was based on multiple differentially expressed genes distributed over multiple branches of amino acid metabolism. A few more genes involved in amino acid catabolism overexpressed in the Selected populations link the aspartate-glutamate metabolism with purines synthesis pathway, which also showed a strong signal of enrichment in differentially expressed genes, and which we examine in more detail below.

### Metabolome analysis points to evolutionary changes in amino acid and purine metabolism

Differences in gene expression between Selected and Control flies suggested that evolutionary adaptation to larval diet affected adult metabolism. We therefore used a broad-scale targeted metabolomics approach to measure the abundance of key polar metabolites involved in multiple core pathways in central metabolism. As for gene expression, the analysis was carried out on the fat body with some adjacent body wall and neural tissue (the “carcass”) of 4-day old females from Selected and Control populations, each raised on either poor or standard diet (additionally, we included a starvation treatment that is analyzed in a later section). After filtering for data quality, we retained 113 metabolites that were quantified (normalized to protein content) in all samples.

The first two principal components extracted from the metabolome data clearly differentiated the Selected versus Control populations, as well as flies raised on poor versus standard diet (Figure 2A; MANOVA on PC1 and PC2 scores, Wilks’ λ = 0.185, *F*_2,9_ = 19.8, *P* = 0.0005 and Wilks’ λ = 0.055, *F*_2,9_ = 77.3, *P* < 0.0001, for regime and diet respectively). Of the 113 metabolites, 57 were found to be differentially abundant between Selected and Control flies at 10% FDR, and also 57 differentially abundant between flies raised on poor versus standard diet (Supplementary file 3). We found no interaction between these two factors either in the univariate analysis (no metabolite passing 10% FDR, lowest *q* = 0.39) or in the PCA (MANOVA, Wilks’ λ = 0.918, *F*_2,9_ = 0.4, *P* = 0.68). Thus, both the evolutionary history of exposure to poor versus standard diet and the within-generation diet treatment had major effects on the metabolome, but, as for gene expression, these effects were largely additive. Therefore, we present the results as a heat map corresponding to effects of these two factors on abundance of individual metabolites (the first two columns of heat maps in Figure 2B; the third column corresponds to the effect of starvation discussed in a later section).

**Figure 2.**
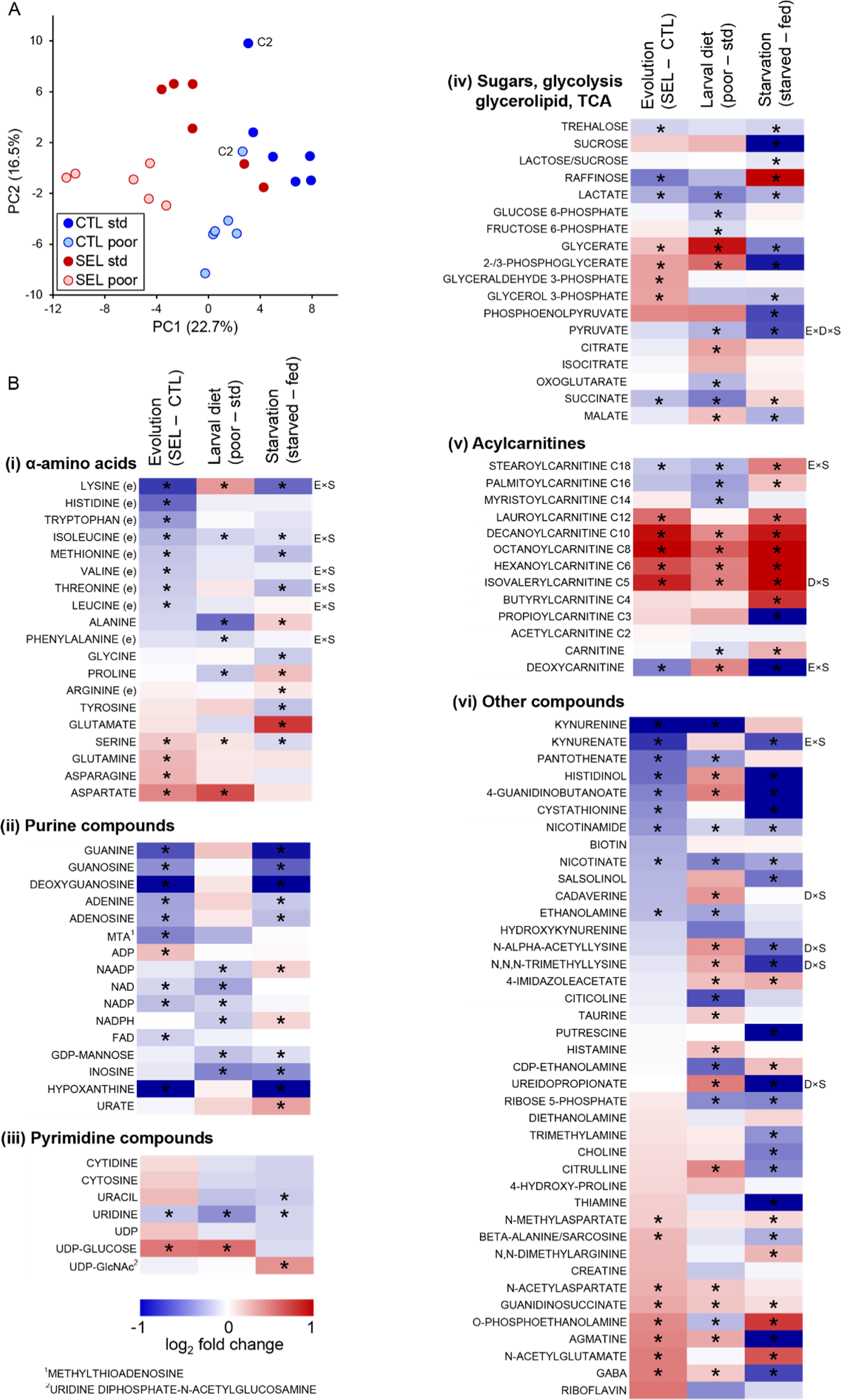
The effect of phenotypic plasticity and evolutionary change on adult metabolite abundance. (A) Principal component score plot based on metabolic signatures composed of 113 robustly quantified metabolites in flies from the six Selected and six Control populations raised on both poor and standard diet (Figure 2—source data 1). The flies were in fed condition. Metabolite abundances for two replicate samples were averaged before the PCA; thus, there is one point per population and diet. The “C2” label indicates control population 2 on either diet; this population deviated from the other Control populations along the PC2 axis; however, we retained it for the univariate analysis. (B) The effects of experimental factors on abundance of individual metabolites. The first two columns of the heat maps indicate, respectively, the effect of evolutionary regime (least square mean difference Selected – Control) and the within-generation (phenotypically plastic) effect of larval diet (least square mean difference poor – standard diet) on the relative concentration of metabolites in fed flies. The third column indicates the main effect of the starvation treatment (least square mean difference between starved and fed flies) on metabolite concentrations in flies from both Selection regimes raised on both diets. Thus red (blue) indicates that a compound is more (less) abundant in Selected, poor diet-raised and starved flies than in Control, standard diet-raised and fed flies, respectively. Asterisks indicate effects significant at *q* < 0.1. Annotations to the right of the heat maps indicate interactions significant at *q* < 0.1: E×S = evolutionary regime × starvation treatment, D×S = larval diet × starvation treatment, E×D×S = three-way interaction; (e) indicates an essential amino acid, C2-C18 for acylcarnitines refers to the length of the acyl chain. For estimates, effects and statistical tests underlying the figure see Supplementary file 3, original data are in Supplementary file 7.

Evolutionary adaptation to larval malnutrition resulted in a striking shift in proteinogenic amino acid concentrations in adult flies: eight essential amino acids were less abundant in the Selected than Control flies, while the reverse was the case for four non-essential ones (Figure 2B(i)). Multiple purine compounds were less abundant in the Selected than Control flies, including both purine nucleobases and nucleosides, and the electron carriers NAD, NADP and FAD; an exception is ADP, which was more abundant in the Selected flies (Figure 2B(ii); ATP was not reliably quantified). No such general trend for reduced abundance was observed for pyrimidines (Figure 2B(iii)), suggesting that the pattern observed for purines is not mediated by changes in synthesis or degradation of nucleic acids.

Selected flies had lower levels of trehalose (the principal sugar circulating in hemolymph) and of lactate than Controls (Figure 2B(iv)), but higher concentrations of compounds linking glycerol with glycolysis, with no indication of differences in concentration of upstream and downstream glycolysis intermediates (Figure 2B(iv)). This suggests changes in fatty acid / triglyceride metabolism. While fatty acids were not included in the targeted metabolome analysis, we did quantify the abundance of acyl-carnitines, i.e., fatty acid residues attached to carnitine, the molecule that transports them in and out of mitochondria. Compared to the Controls, the Selected flies show accumulation of medium chain (C6 – C12) acyl-carnitines (Figure 2B(v)). In contrast, acyl-carnitines with long-chain (C16 – C18) residues, which are typical for insect triglycerides (Stanley-Samuelson, et al. 1988), were less abundant in the Selected than Control flies (Figure 2B(v)).

A number of other metabolites showed differential abundance between Selected and Control flies (Figure 2B(vi)). More than half of them are derivatives of amino acids, including intermediates in synthesis of NAD+ (kynurenine and kynurenate) and cysteine (cystathionine), neurotransmitters (GABA, N-methylaspartate/NMDA and N-acetylaspartate/NAA), and products of amino acid catabolism that do not appear to be functionally linked. Together with increased abundance of isovalerylcarnitine (which carries a residue of deamination of branched-chain amino acids; Figure 2B(v)), these results reinforce the notion that adaptation to the poor larval diet has been associated in wide-ranging changes in amino acid metabolism.

In an attempt to integrate metabolite, gene expression and genome sequence changes associated with adaptation to the poor diet we performed joint pathway analysis in Metaboanalyst v. 5.0 (Xia, et al. 2009). In addition to the metabolites differentially abundant and genes differentially expressed between the Selected and Control populations, we included in this analysis 771 genes previously annotated to single nucleotide polymorphisms (SNPs) differentiated in frequency between these two sets of populations (after 150 generations of experimental evolution; Kawecki, et al. 2021). While most of the nine pathways identified in this analysis were only or mainly enriched in differentially abundant metabolites, two – purine metabolism and alanine, aspartate and glutamate metabolism – involved a mix of metabolites and expression and SNP candidate genes (Supplementary file 4).

We therefore examined the evolutionary changes in these two pathways (imported from the KEGG database; (Kanehisa and Goto 2000)). The alanine-aspartate-glutamate pathway contains multiple links among three of the four amino acids that are overabundant in the Selected flies. However, few of the many enzymes involved in those links showed a signal of differentiation between Selected and Control lines (Figure 3A). In contrast, purine metabolism showed a pattern of upregulation of multiple key enzymes involved in de novo purine synthesis in the Selected population, combined with overabundance of the three amino acids (glutamine, serine and aspartate) that act as donors of four nitrogen atoms that form the basic purine structure (Kastanos, et al. 1997; Salway 2018) and provide (aspartate) a further amino group in the synthesis of AMP from IMP (Figure 3B). Despite this, compared to the Control flies, the Selected flies appeared depleted of nucleosides and nucleobases derived from IMP. We also found that multiple genes involved in AMP-ATP-cAMP conversion were associated with candidate SNPs differentiated in allele frequency between Selected and Control populations. While difficult to interpret functionally, this adds to the evidence that selection driven by the dietary regime targeted purine metabolism.

**Figure 3.**
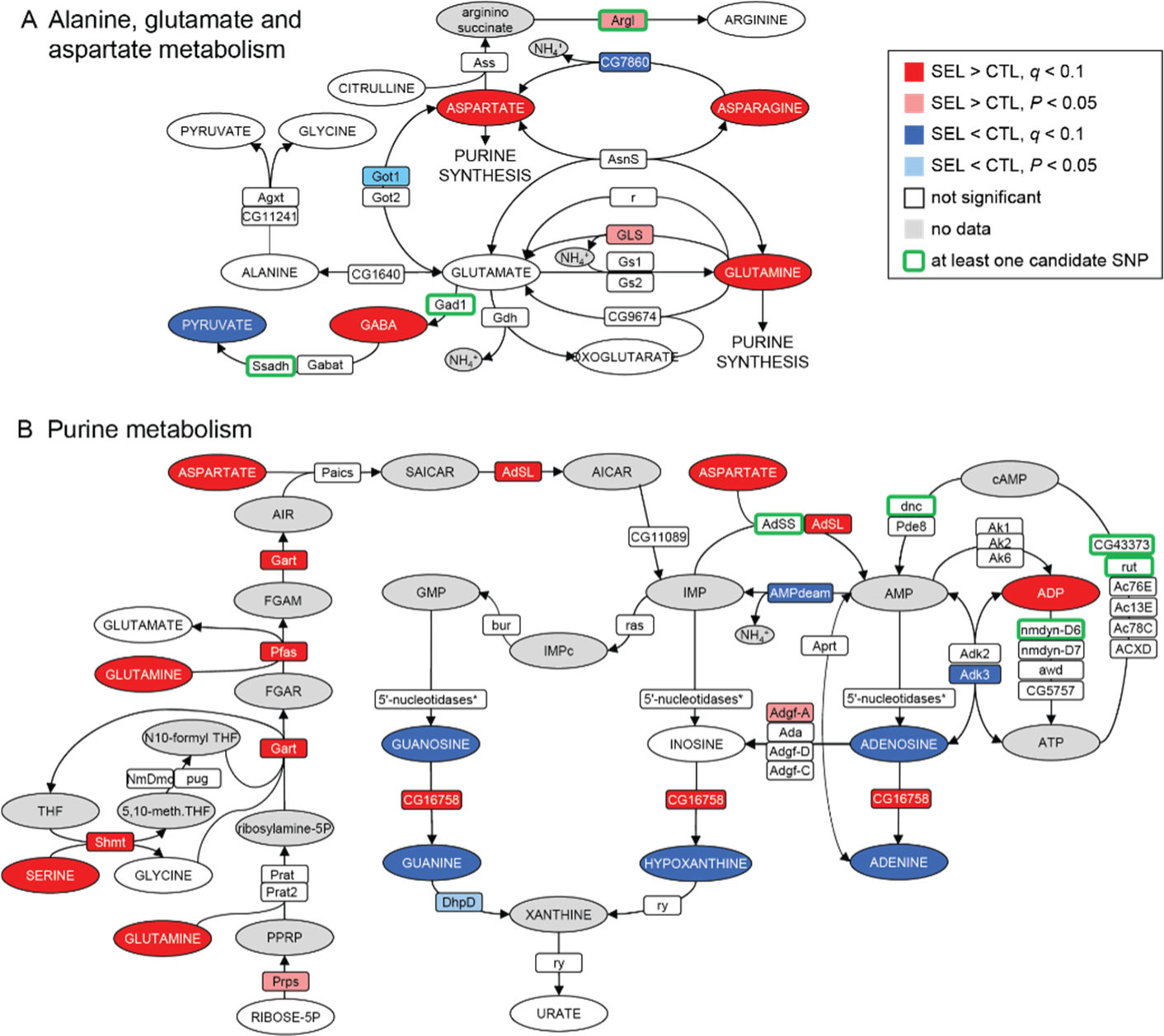
Imprint of evolutionary adaptation to poor larval diet on (A) alanine, glutamate and aspartate metabolism and (B) purine metabolism pathways of adult female flies. Highlighted in color are metabolites differentially abundant and enzymes differentially expressed between the Selected and Control populations, as well as enzymes to which at least one SNP differentiated in frequency between the sets of populations has been annotated. Enzymes differentially expressed at the nominal *P* < 0.05 but not passing the 10% FDR are also indicated. Metabolites in grey were either not targeted, not detected, or the data did not pass quality filtering. The pathways have been downloaded from KEGG (https://www.genome.jp/kegg/pathway.html). For the sake of clarity some substrates and products of depicted reactions that were not differentially abundant or were absent from the data, as well as cofactors, have been omitted. Only genes expressed in female fat body have been included. *5’-nucleosidases: *cN-IIB*, *NT5E-2*, *Nt5B*, *veil*, *CG11883*; none was differentially expressed.

### Plastic response predicts in part evolutionary change of metabolome but not of gene expression

As was the case for Selected versus Control populations, the top GO term enriched for genes differentially expressed in flies raised on the poor versus standard larval diet treatment (corresponding to phenotypic plasticity) was “chitin-based cuticle development” (Figure 1D; Supplementary file 2 table B). This is consistent with differences in body size: flies raised on the poor diet are much smaller than those raised on the standard diet (Kolss, et al. 2009), and thus likely characterized by a greater surface-to-volume ratio and consequently a greater relative surface of cuticle on the carcass. This is, however, the only similarity between the effects of evolutionary regime and of the diet treatment for most enriched GO-terms. No GO terms related to metabolism of amino acids, purines or any other basic metabolic processes were significantly enriched for the effect of diet treatment. Instead, genes differentially affected by the diet treatment were enriched for many terms related to development, reproduction, nervous system and behavior (Figure 1D; Supplementary file 2 table B).

This absence of parallelism between the evolutionary and phenotypically plastic responses to the poor larval diet is also visible at the level of individual genes. Only 42 genes were differentially expressed at 10 % FDR for both of those factors; at least 15 of them are involved in integumentary (i.e., cuticle, epidermis or tracheal) development and most of the remaining ones play a role in the nervous system or transmembrane transport. Of 235 genes that passed the less stringent criterion of being differentially expressed at nominal (not adjusted) *P* < 0.05 for both factors, 123 showed the same sign of expression difference between poor and standard diet as between Selected and Control populations, and 112 showed the opposite sign, not different from what would be expected at random (*P* = 0.13, Fisher’s Exact Test). Across all genes the correlation between the main effect of diet and the main effect of evolutionary regimes was weakly negative (*r* = −0.22, *P* < 0.0001, *N* = 8701; Figure 4A), as was the corresponding correlation for genes annotated to metabolism (GO metabolic process, *r* = −0.26, *P* < 0.0001, *N* = 4606). Thus, except for genes involved in the cuticle maturation, the plastic (within-generation) response of adult gene expression pattern to larval undernutrition did not in general predict the evolutionary, genetically-based response. If anything, evolution tended on average to oppose phenotypically plastic responses of gene expression to larval diet.

**Figure 4.**
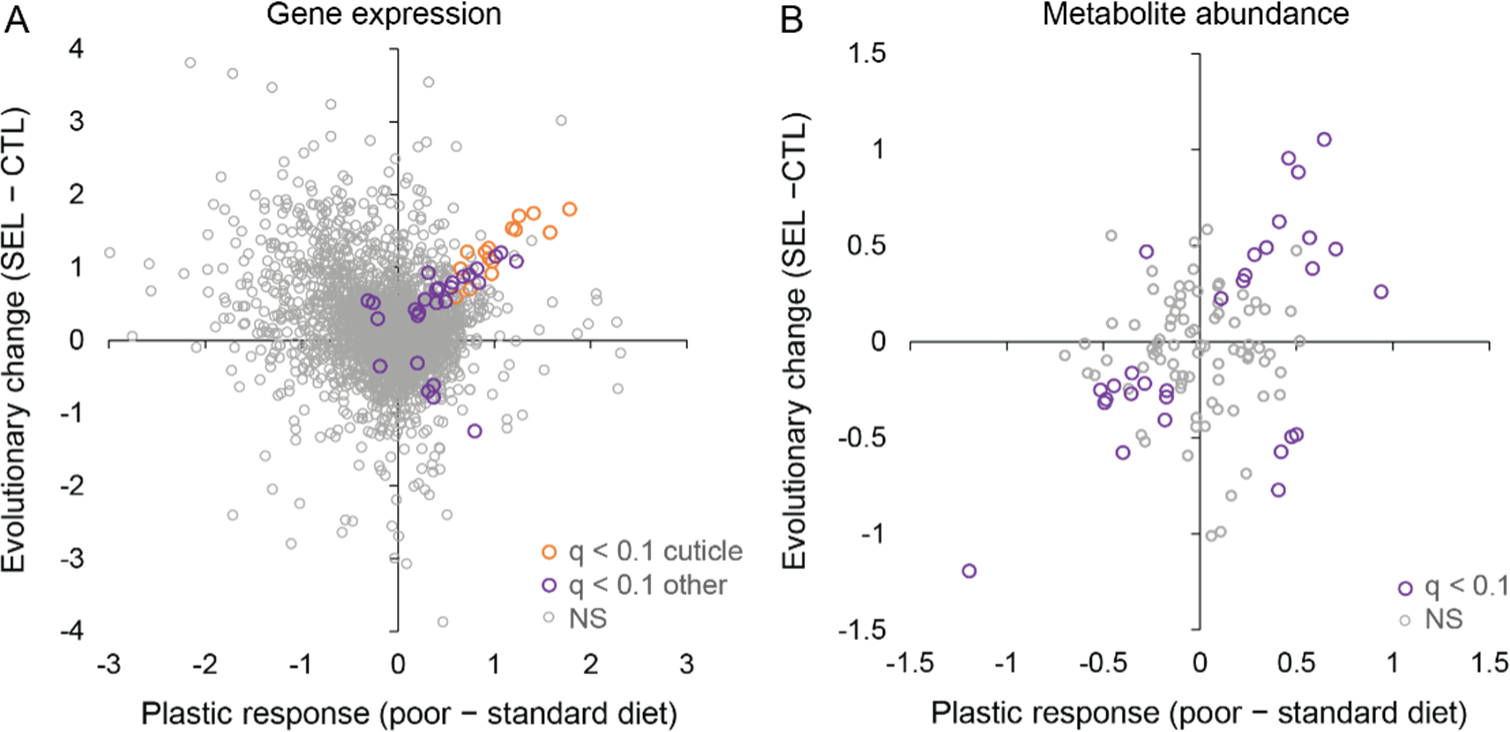
Relationship between phenotypic plasticity and genetically-based evolutionary change of adult gene expression (A) and metabolite abundance (B) in response to larval diet. The plastic responses and evolutionary change for each gene and metabolite were estimated, respectively, as the main effect of larval diet and evolutionary regime in the factorial statistical models (see Methods). Color highlights genes and metabolites with significant effects (*q* < 0.1) for both diet and regime; in (A) orange symbols indicate significant genes involved in cuticle, integument or tracheal development (Figure 4—source data 1).

By contrast, the evolutionary change of metabolome driven by larval diet did to some degree parallel the phenotypically plastic response (correlation across all metabolites *r* = 0.32, *P* = 0.0005, *N* = 113; Figure 4B). Of the 30 metabolites that were significant at 10% FDR for both evolutionary regime and diet treatment, 25 showed the same sign of the difference between Selected and Control flies as the difference between flies raised on the poor vs standard diet (Fisher’s Exact Test, *P* = 0.0005). In particular, compared to flies raised on standard diet, flies raised on the poor diet showed a lower abundance of long-chain acyl-carnitines but an excess of medium-chain ones (Figure 2B(v), middle column), and a lower abundance of NAD+ and NADP (Figure 2B(ii), middle column) and their precursors nicotinate and kynurenine (Figure 2B(vi), middle column), paralleling the patterns in Selected relative to Control flies (the left column of the respective heat maps). Some of the plastic effects of the poor diet treatment on sugar metabolism were also similar to those of evolutionary response, notably an underabundance of lactate and succinate, and an overabundance of UDP-glucose and of 2-/3-phosphoglycerate and glycerate, the latter accentuated by the underabundance of upstream and downstream glycolysis intermediates (Figure 2B(iv)).

More than a dozen modified amino acids and products of amino acid degradation were also affected, but these changes were not correlated with corresponding differences between the Selected and Control flies. Although several proteinogenic amino acids showed differential abundance depending on the diet treatment, these differences were also quite different from those due to the evolutionary regime and did not seem to show a consistent relationship with the properties of the amino acids. Finally, with the exception of inosine, the plastic response to diet did not detectably affect the abundance of purine nucleobases and nucleotides that do not contain a nicotinamide group (Figure 2B(ii)). These results are consistent with the absence of signal of an effect of the diet treatment on amino acid and purine metabolism in gene expression patterns presented above.

### Much of evolutionary change in gene expression and metabolome is conserved between life stages

The Selected and Control populations evolved under different larval diets but the adult diet they experienced was the same. Hence, while the divergence between them in larval gene expression and metabolite abundance is presumably driven by immediate nutrient availability, selection at the adult stage would likely result from the need to compensate for carry-over effects of larval diet. It does not seem likely that these two selection pressures would favor broadly similar changes in gene expression and metabolism. However, such a broad similarity would still be expected if the genetic variants favored by selection on the larvae had similar effects across the stages. We therefore explored whether the divergence between Selected and Control populations in gene expression and metabolite abundance was correlated between the two life stages. To do so we combined the present data from adult carcasses with previously published RNAseq and targeted metabolome data from whole 3rd instar larvae, collected at generation 190 (Erkosar, et al. 2017) and generation 264 (Cavigliasso, et al. 2023), respectively.

Of 424 genes that were differentially expressed at nominal *P* < 0.05 between Selected and Controls at both adults and larvae (Supplementary file 5), 377 (89%) showed the same direction of change on poor diet, more than expected at random (Fisher’s Exact Test, *P* < 0.0001, Figure 5A). Eighty-four genes passed the more stringent criterion of 10 % FDR at both stages; of those, 78 (93%) showed the same direction of change in the two stages (Fisher’s Exact Test, *P* < 0.0001). The estimates of log-fold change were even correlated across all 8437 genes shared between the data sets (*r* = 0.35, *P* < 0.0001; Figure 5A). Thus, many of the differences between Selected and Control populations in the adult gene expression patterns paralleled those observed in the larvae.

**Figure 5.**
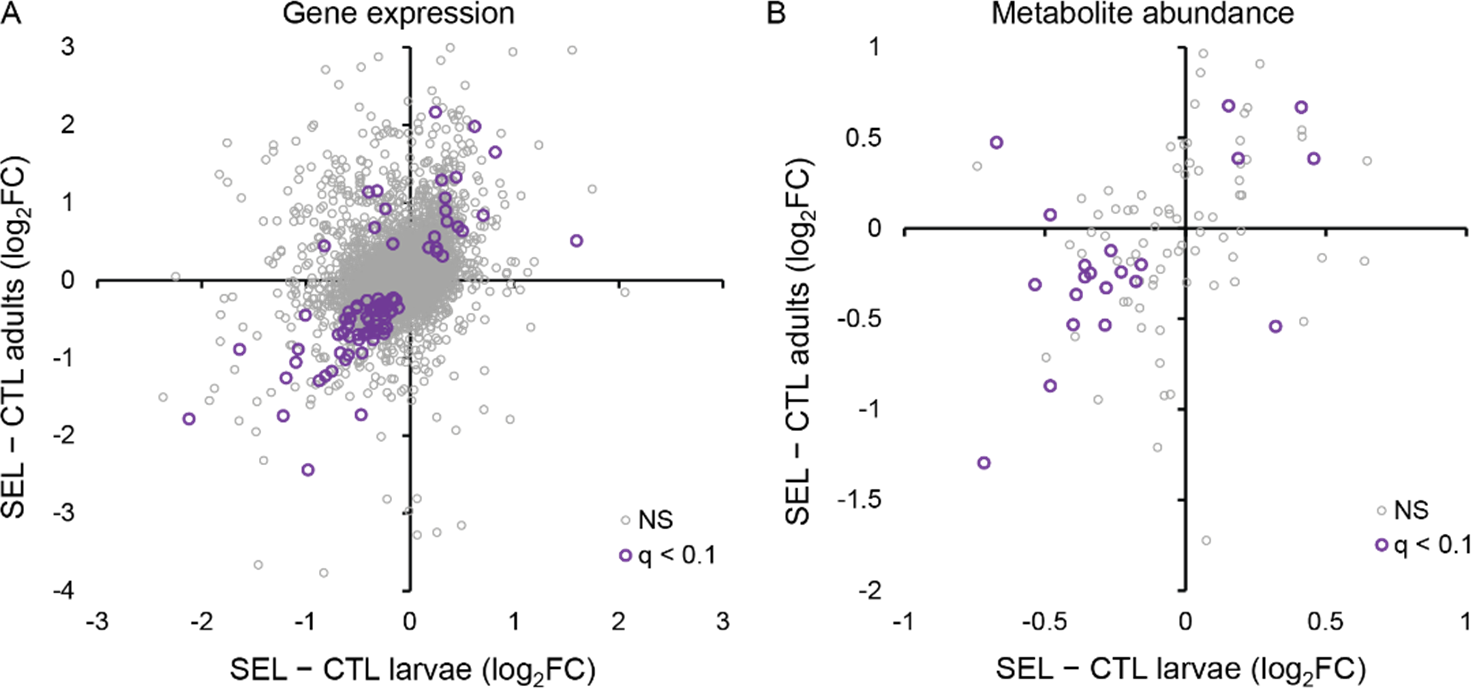
The relationship between the evolutionary change (i.e., the difference between the Selected and Control populations) in gene expression and metabolite abundance at the larval and adult stage. (A) Correlation of evolutionary changes in gene expression (all genes: *r* = 0.35, *N* = 8437, *P* < 0.0001; genes with *q* < 0.1: *r* = 0.74, *N* = 84, *P* < 0.0001). Five genes (not passing *q* < 0.1) are outside of the range of the plot. (B) Correlation of evolutionary changes in metabolite abundance (all metabolites: *r* = 0.37, *N* = 97, *P* = 0.0002; metabolites with *q* < 0.1: *r* = 0.55, *N* = 21, *P* = 0.0091). The estimates plotted are for larvae and adults raised on the poor larval diet (see Methods); *q* < 0.1 refers to genes or metabolites that pass the 10% FDR threshold for both stages; NS are the remaining genes or metabolites (Figure 5—source data 1, sheet A and B).

Genes that were differentially expressed in Selected versus Control population in both adults and larvae were enriched in several terms related to transmembrane transport and iron homeostasis (Supplementary file 2 table D). Differences in gene expression of the larvae showed enrichment in lipid and carboxylic acid metabolism, as well as in a number of GO terms linked to cell proliferation and development, including that of the nervous system (Supplementary file 2 table C). The larval differentially expressed genes were not enriched in GO terms linked to amino acid or purine metabolism. However, 18 out of 99 genes in GO term “alpha amino acid metabolic process” and 16 out of 107 in GO “purine-containing compound biosynthetic process” were significantly different between Selected and Control larvae at 10% FDR (Supplementary file 6). Furthermore, the differences between Selected and Control populations across all genes in those GO terms were positively correlated between larvae and adults (amino acid metabolism: *r* = 0.40, *P* < 0.0001, *N* = 94; GO purine synthesis: *r* = 0.47, *P* < 0.0001, *N* = 74). Thus, while the amino acid metabolism and purine synthesis do not show disproportionate changes in gene expression in the larvae compared to other GO terms, the expression of multiple genes in those pathways has clearly been affected in a similar way as observed in adults.

Similarity between the effect of experimental evolution on larvae and adults was also apparent for the metabolome. Across all metabolites, the difference in metabolite abundance between the Selected and Control populations in the adults was positively correlated with the analogous difference in the larvae (*r* = 0.37, *P* = 0.0002; Figure 5B). This correlation was even more pronounced across the 21 metabolites that were differentially abundant at both stages (*r* = 0.55, *P* = 0.009); of those, 18 showed the same direction of change in larvae and adults, significantly more than expected by chance (Fisher’s Exact Test, *P* = 0.011).

In contrast, in the same data set we detected no correlation between the direct (phenotypically plastic) responses of larval and adult metabolome to larval diet, whether across all metabolites (*r* = 0.01, *P* = 0.89) or across the 35 metabolites detected as differentially abundant in response to diet at both stages (*r* = 0.17, *P* = 0.34; Figure 5 – figure supplement 1). Of that last group, 19 showed the same sign of change at both stages while 16 showed opposite signs, not different from random expectation (*P* = 0.73). This suggests that differences in metabolite abundance accrued at the larval can be erased within a few days of adult life. Hence, differences between Selected and Control adults in metabolite abundance are more likely to be mediated by differences in adult gene expression than by differential accumulation of metabolites during the larval stage.

### Metabolic profile of Selected flies tends towards starved-like state

In addition to assessing the metabolome of flies directly sampled from the (standard) adult diet, we also assessed the metabolome of Selected and Control flies after they have been subject to 24 h of total food deprivation (moisture was provided). The effect of the starvation treatment on the metabolic phenotype is also a phenotypically plastic response – to a different form of nutritional stress and one applied to adults rather than larvae. Our motivation was twofold. First, a comparison of fed and starved flies would reveal how the metabolome changes under an ongoing nutritional stress, and thus potentially help interpret metabolome changes driven by genetically-based adaptation to recurrent nutrient shortage at the larval stage (i.e., the effect of evolutionary regime). Second, adult flies from Selected populations are less resistant to starvation than Controls (Kawecki, et al. 2021). Hence, we expected that the metabolome of the Selected and Control populations might throw some light on the mechanisms underlying these differences in starvation resistance.

As expected, 24 h of starvation had a strong effect on fly metabolome. Principal component analysis cleanly separated starved from fed samples along the first PC axis (Figure 6A), while larval diet mainly defined the second PC axis. Seventy-four out of the 113 metabolites were detected to be differentially abundant (at 10% FDR) between fed and starved flies (in terms of the main effect of starvation treatment in the full factorial model, Supplementary file 3). Of 39 metabolites affected by both starvation treatment and the evolutionary regime, 29 showed the same direction of change between fed and starved flies as between Control and Selected flies, significantly more than expected by chance (*P* = 0.006, Fisher’s Exact Test). In particular, in parallel to Selected flies relative to Controls, starved flies showed a decrease in multiple purine metabolites and two forms of vitamin B3 (nicotinate and nicotinamide), and an increase in medium-chain acyl-carnitines (Figure 2B). For some other metabolites the effect of starvation differed markedly from that of the evolutionary regime. In particular, several intermediates of the glycolysis pathway were markedly less abundant in starved than fed flies (Figure 2B) whereas the surplus of acyl-carnitines also applied to those with long acyl chains. This possibly reflects a shift to catabolizing lipid stores for ATP generation following the depletion of carbohydrates.

**Figure 6.**
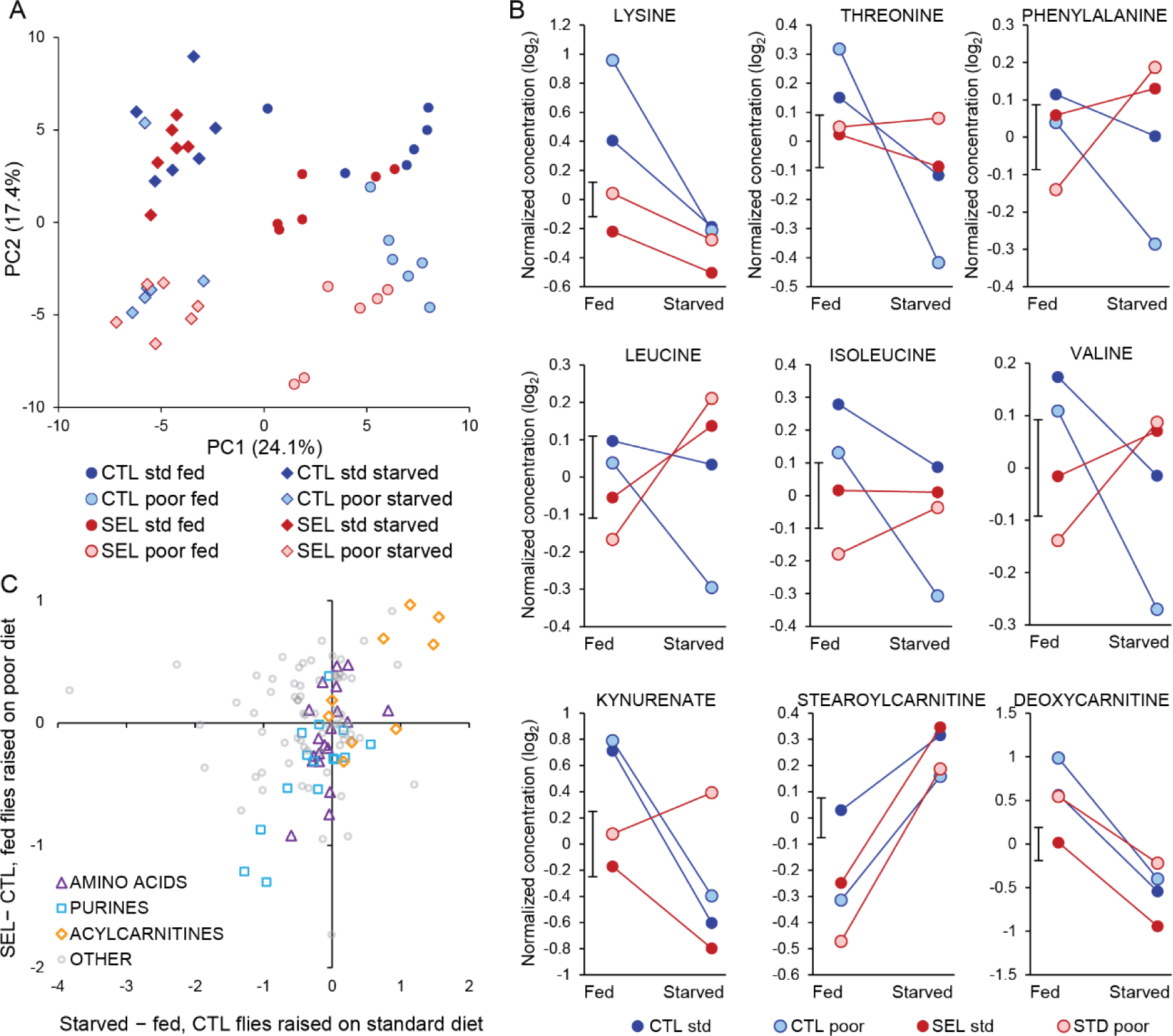
(A) Principal component score plot on metabolite abundance for all conditions (Selected vs Control, poor vs standard larval diet, fed vs starved flies). The two replicate samples from the same population × diet × starvation treatment combination were averaged (Figure 6—source data 1). (B) Metabolites that show significant (*q* < 0.1) interaction between the evolutionary regime and starvation treatment. Plotted are least-square means obtained from the full factorial general mixed model (Figure 6—source data 2). The bar next to the Y-axis indicates ± standard error of the least square means (i.e., its width corresponds to 2 SEM); the standard error is identical for all treatments because the least square means were estimated from the general mixed model and the design is balanced. For compounds showing other interactions see Figure 6—figure supplement 1 (C) The relationship between the effect of 24 h starvation (starved minus fed flies) on metabolite abundance in Control flies raised on the standard diet and the evolutionary change (Selected minus Control) quantified in fed flies raised on poor diet (Figure 6—source data 3). Three categories of metabolites of interest are highlighted; “acylcarnitines” only include those with even-numbered acyl chain (resulting from catabolism of triglycerides) but not those with 3C and 5C chains (which are products of amino acid catabolism). Pearson’s correlations: amino acids *r* = 0.50, *N* = 19, *P* = 0.030; purine compounds *r* = 0.76, *N* = 16, *P* = 0.0006; acylcarnitines *r* = 0.74, *N* = 9, *P* = 0.022; other metabolites *r* = 0.02.

The pattern for proteogenic amino acids also appeared rather different (Figure 2B). However, the focus on the main effects of starvation treatment on amino acid abundance is somewhat misleading because for six of the ten essential amino acids the response to starvation differed between the Selected and Control flies. For all six the sign of the interaction was the same: while the levels of all these amino acids decreased in Control flies when they starved (*q* < 0.1), their abundance declined less (lysine), remained similar (threonine, isoleucine) or even increased (phenylalanine, leucine, valine; *q* < 0.1) in starving Selected flies (Figure 6B). The other three metabolites with significant regime × starvation interaction showed different patterns (Figure 6B). There were also six metabolites with a significant larval diet × starvation or an interaction among all three experimental factors; they did not appear to be functionally connected or to share a common pattern (Fig. 6—figure supplement 1).

These results imply that evolutionary adaptation of the Selected populations to larval undernutrition had consequences for the way their essential amino acid metabolism responds to food deprivation at the adult stage. Furthermore, even though in a normal fed state the Selected flies showed lower abundance of most essential amino acids, they could maintain or increase their levels during 24 h of starvation. This suggests that their low concentrations in the fed state are not due to their absolute shortage but to differential allocation and use. In total, abundance of seven essential amino acids declined significantly upon starvation in Control flies, making the parallel with the difference between Selected and Control flies in fed state more evident. Indeed, the evolved differences in abundance of all proteogenic amino acids between Selected and Control flies were significantly positively correlated with the presumably ancestral response of standard diet-raised Control to starvation; the same was the case for purine compounds and acyl carnitines (Figure 6C).

Differential response of the Selected and Control populations to starvation is supported by regime × starvation interaction on PC1 scores (*F*_1,10_ = 10.7, *P* = 0.0084) – the metabolic state of Selected flies changed less upon starvation than that of Controls. This does not imply that the metabolome of Selected flies was more robust to starvation. Rather, fed flies from the Selected populations were situated closer to starved flies along the starvation-loaded PC1 axis than fed Control flies (Figure 6A; *F*_1,17.8_ = 14.0, *P* = 0.0015, GMM on PC1 scores); in the starved condition they converged to a similar state (*F*_1,17.8_ = 0.0, *P* = 0.99; see Supplementary file 7 for detailed statistics). Overall, the comparison of the (plastic) response of metabolome to starvation indicates that prominent aspects of the adult metabolic profile of Selected populations evolved in the direction resembling effects of starvation.

### Adult cost of larval adaptation

The above results demonstrate that evolutionary adaptation to nutrient-poor larval diet has led to significant changes in adult metabolism. It is not clear, however, to what degree these changes may have evolved to improve fitness of adults that experienced poor diet as larvae, rather than being carry-over effects or trade-offs of changes driven by selection acting on the larval stage. If the former, adults from the Selected populations should perform better than those from the Control populations when both are raised under the conditions of the poor larval diet regime. If the latter, the fitness of Selected adults raised on poor diet should be similar to or even lower than the fitness of Control adults developed on the same poor diet.

To test these alternative predictions, we compared the fecundity of females from Selected and Control populations raised on the poor larval diet, transferred to standard diet after eclosion, supplemented with ad-libitum live yeast for 24 h before oviposition, and allowed to oviposit overnight. These conditions mimicked those under which Selected populations have been maintained and propagated in the course of their experimental evolution; a female that laid more eggs in this time window would contribute proportionally more offspring to the next generation. The rate at which females can convert nutrients into eggs within this short time window is thus arguably a key measure of their fitness, and is inherently dependent on the metabolism of nutrients, in particular proteins, lipids and nucleic acids. Despite having evolved under these conditions, Selected females laid only about 2/3 of the number of eggs produced by the Control females, whose ancestors did not face the poor larval diet (Figure 7A). Fly ovaries consist of a variable number of branches called ovarioles; in each ovariole eggs are produced sequentially one by one. The rate of egg production may thus be limited by the number of ovarioles (Schmidt, et al. 2005), which is determined during larval development (Bergland, et al. 2008). Indeed, we found that poor diet-raised females of the Selected populations tended to have slightly fewer ovarioles than their Control counterparts (Figure 7B). However, even adjusted for the difference in ovariole number, the Selected flies produced substantially fewer eggs than Controls (Figure 7C).

**Figure 7.**
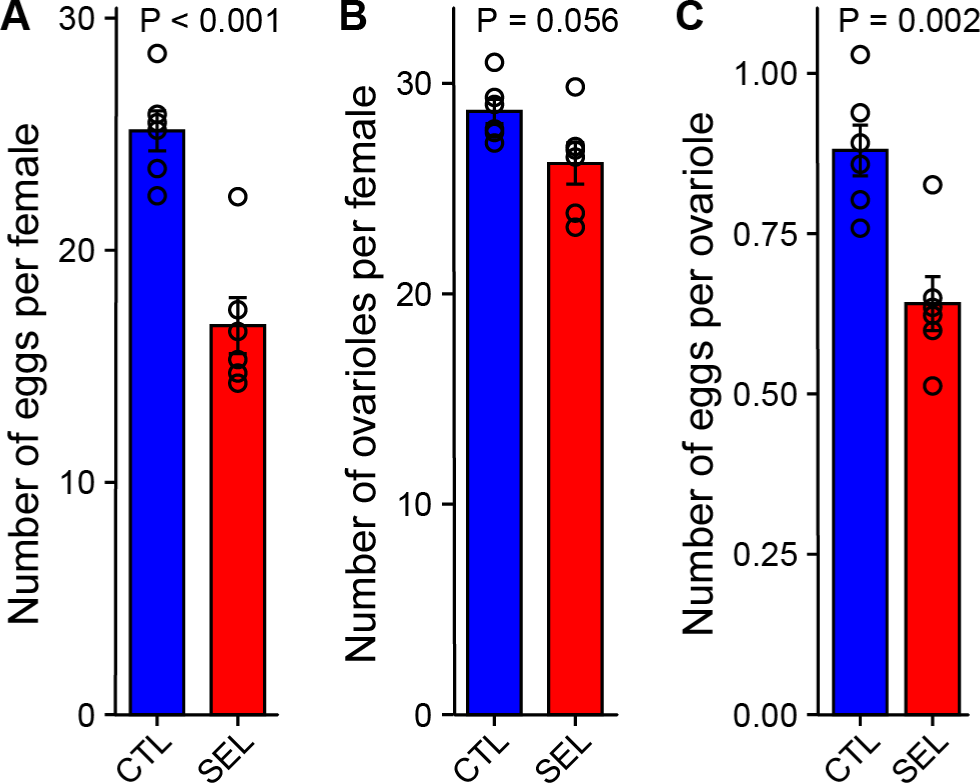
Selected populations pay a fecundity cost for larval adaptation to poor diet. (A) The number of eggs laid overnight by females from the Selected and Control (GMM; *F*_1,8_ = 44.5, *P* = 0.0002, *N* = 16 per evolutionary regime, 2-3 per replicate population). (B) The number of ovarioles (left + right ovary) per female (GMM; *F*_1,10_ = 4.7, *P* = 0.056, *N* = 72 per regime, 6 per replicate population). (C) The mean number of eggs per ovariole (one-way ANOVA; *F*_1,10_ = 17.1, *P* = 0.002, *N* = 6 populations per regime). The females experienced the dietary conditions, under which the Selected populations evolved: the larvae were raised on the poor diet; the adults were transferred to the standard diet and additionally provided live yeast 24 h before oviposition. Bars are means, error bars correspond to ± SEM, the symbols to means of replicate populations (Figure 7-source data 1).

*Drosophila* are “income breeders” in that they directly convert acquired nutrients into eggs rather than relying on previously accumulated reserves (O’Brien, et al. 2008). The fecundity difference thus implies that the Selected populations are less effective than Controls at converting dietary nutrients into the nutritional and/or generative components of the egg even when raised under the conditions under which the former but not the latter evolved during more than 200 generations.

## Discussion

### Costly correlated evolution of adult metabolism?

Experimental evolution under chronic larval undernutrition resulted in major shifts in gene expression patterns and metabolite abundance at the adult stage, replicable across six independent experimental (Selected) populations of *Drosophila melanogaster*. This occurred even though adults of all populations experienced standard diet in the course of the experimental evolution. These evolved differences in adult gene expression and metabolite abundance between Selected and Control populations were broadly positively correlated with the corresponding differences between Selected and Control larvae. This was the case even though expression and metabolites were quantified in whole body in larvae but in dissected carcasses in adults, and despite these assays being performed dozens of fly generations apart. Therefore, the observed correlations between adult and larval evolutionary responses may have underestimated their actual similarity.

Even though adults of both Selected and Control populations experienced the same standard diet, they might still have been subject to differential selection on their adult metabolism. In particular, selection on Selected adults may have favored changes that compensate for lingering consequences nutritional hardship endured during the larval stage. However, it seems difficult to conceive why such compensatory responses of a well-fed adult should be mediated by similar physiological adjustments as those that promote larval growth under severe undernutrition. A more plausible explanation for the similarity in evolutionary changes between larvae and adults is that most genetic variants available for evolution affected gene expression of larvae and adults similarly. As a consequence, many of the adult gene expression changes would represent correlated responses to selection on the larval stage rather than the direct response to selection on adult performance (Lande and Arnold 1983).

Such correlated responses may be detrimental to performance of one or both life stages (Collet and Fellous 2019). Consistent with this prediction, when raised on the poor diet and transferred to the standard diet as adults, the Selected flies were less effective than Controls in converting dietary nutrients into eggs, even though these were the conditions under which the former had evolved and to which the latter were exposed for the first time. (An analogous result on fecundity has been reported in another evolution experiment (May, et al. 2019).) We have previously reported a lower fecundity of Selected females compared to Controls when both were raised on standard diet (Kolss, et al. 2009); we interpreted that finding as a manifestation of reduced performance of Selected populations when raised on the standard larval diet. The present results imply that Selected adults perform less well than Control adults irrespective of the larval diet on which they have been raised. Other things being equal, a female’s contribution to the next generation under the experimental evolution regime is proportional to the number of eggs she lays within a time window similar to that used to quantify fecundity. Thus, the fecundity reduction we found in Selected adults represents a significant fitness trade-off of the improved ability of the Selected larvae to survive and develop under extreme nutrient shortage.

### Amino acid catabolism and starved-like metabolic profile?

While Selected and Control adults differed in abundance of a variety of metabolites, a most striking pattern was reduced abundance of eight out of ten essential amino acids in Selected flies. Fecundity in *Drosophila* is limited by essential amino acids (Grandison, et al. 2009), in particular those in short supply relative to the needs of fly protein synthesis, which for a yeast-based diet are methionine and leucine (Piper, et al. 2017). In addition, several essential amino acids (methionine, valine, isoleucine and in particular leucine, all less abundant in Selected flies) act as signaling molecules that promote protein synthesis by activating TOR complex 1 (Wolfson, et al. 2016; Antikainen, et al. 2017; Gu, et al. 2022). Most of protein synthesis activity in adult female fat bodies is directed towards synthesizing egg proteins (Piper, et al. 2017; Gupta, et al. 2022). Thus, while we have not demonstrated a direct causal link, lower abundance of most essential amino acids in the Selected flies is consistent with their lower fecundity.

A reduction in free amino acid abundance could result from a lowered supply from nutrition and/or from a higher rate of their use for protein synthesis. However, these mechanisms should lead to a general depletion of free amino acids, which is not what we observed. Rather, in contrast to essential amino acids, several non-essential amino acids were overabundant in Selected flies; these overabundant amino acids are not enriched in dietary yeast relative to their need for *Drosophila* protein synthesis (Piper, et al. 2017). Thus, rather than being explained by differences in dietary amino acid acquisition or use in protein synthesis, the differences in amino acid abundance between Selected and Control populations appear consistent with differential use of amino acids in metabolism.

As adults, larvae of Selected populations also show lower abundances than Control larvae of multiple amino acids, including six essential, and a higher concentration of uric acid (Cavigliasso, et al. 2023), an end product of purine metabolism and the main form in which nitrogen is excreted in insects (Salway 2018; Cohen, et al. 2020). This suggests that Selected larvae catabolize amino-acids and excrete nitrogenous waste products at a higher rate, a hypothesis further supported by their increased accumulation the heavy isotope of nitrogen ^15^N (Cavigliasso, et al. 2023). Although we found no difference in the levels of uric acid (urate) in the adult metabolome, it should be kept in mind that the adult metabolome was quantified carcasses, which only include fragments of Malpighian tubules, the excretory organs in which uric acid is synthesized (Cohen, et al. 2020). It is notable that Selected flies show overabundance of the three amino acids – glutamine, aspartate and serine – that contribute nitrogen atoms to purine / uric acid synthesis pathway, as well as overexpression of multiple genes involved in that process. Thus, while we do not have direct evidence for this, it remains possible that Selected flies catabolize amino acids and excrete uric acid at a higher rate than Controls also at the adult stage.

Increased catabolism of amino acids – sourced from autophagy – is one of the hallmarks of the physiological response to starvation (Scott, et al. 2004). We found that several differences in metabolome between Selected and Control flies resemble the effects of starvation. In addition to reduction in abundance of several essential amino acids, this includes lower concentration of purine nucleotides and nucleosides, lower levels of trehalose and lactate, and higher abundance of acylcarnitines hinting at increased catabolism of fatty acids. Thus, the metabolic profile of Selected flies, in their normal fed and reproductively active state, resembles that of flies that have been starving. Consistent with the link with starvation, the genomic architecture of differentiation between the Selected and Control populations includes many candidate genes for starvation resistance (Kawecki, et al. 2021). It is tempting to speculate that, as a byproduct of genetic adaptation to poor larval diet, the Selected populations have become programmed to express a starved-like adult metabolic phenotype, which might explain their low fecundity.

### Relationship between phenotypic plasticity and evolutionary change

The evolutionary changes in gene expression did not in general recapitulate the phenotypically plastic responses of adult expression to larval diet. The abundance of a subset of metabolites did evolve in the direction that mimicked the plastic response, but this was not the case for most of the other metabolites. Congruence between the directions of phenotypically plastic response and evolutionary change driven by the same environmental factor has been interpreted as evidence for adaptive nature of the plastic response; conversely, where evolutionary change went in the opposite direction to the plastic response, the latter has been deemed maladaptive (Yampolsky, et al. 2012; Ghalambor, et al. 2015; Huang and Agrawal 2016; Josephs, et al. 2021). We believe this interpretation is not applicable to our results. First, as we argued elsewhere ((Cavigliasso, et al. 2023)), a plastic response to a novel environment may be adaptive in terms of direction but overshoot the optimum phenotype; in such a case, an evolutionary change in the opposite direction would be favored despite the initial plastic response being adaptive. Second, the above interpretation assumes that evolutionary changes are mostly adaptive. However, as we have argued above, the evolutionary responses of adult gene expression and metabolism to larval undernutrition are likely to a large degree maladaptive costs of physiological adaptations favored at the larval stage. Thus, even plastic responses optimal from the viewpoint of adult fitness may have been reversed by the evolutionary change. These considerations imply that assessing the adaptive (or otherwise) nature of phenotypically plastic responses based on the direction of the evolutionary change may be misleading. Furthermore, our experimental results contradict the often made assertion that phenotypic plasticity “drives” evolutionary change (Baldwin 1896; Pigliucci and Murren 2003; Moczek, et al. 2011; Laland, et al. 2015).

### Evolutionary constraints on regulatory flexibility?

Like in all holometabolous insects, most larval tissues in *Drosophila* disintegrate during metamorphosis and the rest (e.g., brain) undergo extensive remodeling. Most of adult structures and organs are formed de novo from progenitor cells, resulting in an adult that is very different morphologically from the larvae, and physiologically specialized for a different function (reproduction rather than growth). In particular, cells of the larval fat body dissociate from one another and undergo autophagy and apoptosis during metamorphosis; the adult fat body develops anew from adult progenitor cells although possibly including some remaining larval fat body cells (Li, et al. 2019). A major advantage of this complex development is thought to be that it decouples larval and adult gene expression, promoting independent evolution of larval and adult phenotypes (Moran 1994; Rolff, et al. 2019). Contrary to this notion, our results suggest that, at least on the scale of hundreds of generations, evolutionary changes in physiology driven by selection acting on larvae may have maladaptive pleiotropic effects on adult physiology. If this is the case in a holometabolous insect, such evolutionary non-independence of juvenile and adult physiology would likely be more pronounced in species with more developmental continuity between juvenile and adult tissues and organs.

Our study is thus relevant to understanding of constrains on the evolutionary refinement of physiology and life history of metazoans. Complex multicellularity crucially depends on ability of the genome to express its genes differently in different cell types and life stages. The existence of specialized cells and organs that express greatly different yet highly coordinated and functional metabolic phenotypes from the same genome testifies to the power of regulatory evolution. On the other hand, there has been increased recognition that the ability of evolution to independently shape phenotypes of different life stages and sexes may be significantly constrained. Such constrains are expected to emerge from the complexity of gene regulatory and metabolic networks (Wagner 2011; Sorrells, et al. 2015; Schaerli, et al. 2018). One manifestation of such constraints is “intralocus sexual conflict” (sexually antagonistic pleiotropy), whereby simultaneous optimization of female and male phenotypes is hindered by constraints on independent evolution of gene expression in the two sexes (Rice 1984; Pischedda and Chippindale 2006; Hollis, et al. 2014; Veltsos, et al. 2017). Similarly, the developmental theory of aging postulates that gene expression and metabolism are optimized for maximizing performance at a young age and fail to adjust in later age in ways that could improve reproductive lifespan or healthspan (de Magalhaes 2012; Gems and Partridge 2013), an idea increasingly supported by experimental data (Carlsson, et al. 2021). This apparent metabolic inertia of aging individuals might be explained by selection at old age being weak (Medawar 1952; Hamilton 1966; Partridge and Barton 1993). However, our results suggest similar evolutionary constraints linking the metabolism of juveniles and young adults in their reproductive prime, before the age-related decline in the strength of natural selection sets in (Hamilton 1966). Such constraints would hinder evolutionary optimization of juvenile and adult gene expression and metabolism if optima differ between the stages (Collet and Fellous 2019).

## Materials and Methods

### Diets, experimental evolution and fly husbandry

Two diets were used for this study. The “standard” diet consisted of 12.5 g dry brewer’s yeast, 30 g sucrose, 60 g glucose, 50 g cornmeal, 0.5 g CaCl2, 0.5 g MgSO4, 10 ml 10% Nipagin, 6 ml propionic acid, 20 ml ethanol and 15 g of agar per liter of water. The “poor” diet contained 1/4 of the concentrations of yeast, sugars and cornmeal, but the same concentrations of the other ingredients. All experiments were carried out at 25°C and 12:12 h LD cycle.

Six Selected and six Control populations were all derived from the same base population originally collected in 1999 in Basel (Switzerland) and maintained on the standard food for several years before the evolution experiment started in 2005 (Kolss, et al. 2009). The six replicate populations per evolutionary regime constitute the main units of replication in this study; their number was limited by workload considerations. By the time of the experiments reported in this paper the Selected populations had been maintained on the poor larval diet for over 230 generations; the Control populations were maintained in parallel on the standard larval diet. Larval density was controlled at 200-250 eggs per bottle with 40 ml of food medium. Adults of both regimes were transferred to standard diet within 14 days of eggs laying (sometimes a day or two later when not enough adults have emerged from the poor diet) and additionally fed live yeast 24 h before egg collection to stimulate egg production. The target adult population size was 180-200 individuals.

Flies used in the experiments reported here were raised using similar procedures. Prior to all experiments reported in this study all populations were raised on the standard diet for at least two generations to minimize the effects of maternal environment (Vijendravarma, et al. 2010). For RNAseq and metabolome analyses flies of both Selected and Control populations were raised on both larval diets. Eggs to establish the next generation were collected by allowing flies to oviposit overnight on orange juice agar sprinkled with live yeast. During the egg collection, the eggs were washed with water to remove any traces of diet at the surface – a procedure that also removes much of microbiota. To control the colonization by microbiota, eggs were re-inoculated using parental feces from a mix of adult flies from all 12 populations. These adults were left in a petri dish and a wedge of food for 48 h; they and the food were subsequently removed, the feces were washed from the surfaces of the petri dish with PBS, filtered to remove eggs and debris, and adjusted to OD_600_ = 0.5. Each larval culture was established with 200 eggs transferred to a bottle with 40 ml of poor or standard diet, with 300 µl of the feces suspension pipetted on top; based on plating, this inoculum contains about 10^3^ CFU. All experiments were performed on females, aged 4-6 days old from eclosion. Selected larvae develop on the poor diet faster than Controls (Kolss, et al. 2009; Erkosar, et al. 2017). To ensure that the females from Selected and Control populations can be collected synchronously, we initiated the larval cultures in a staggered manner over several days, so that we could collect females from around peak of emergence and at the same time for Selected and Control populations despite the difference in developmental time. For RNAseq and metabolome quantification, adults of both sexes were collected around the peak of emergence, transferred to fresh standard diet and allowed to mate freely for three days before females were collected. For fecundity and ovariole measurements we collected virgin females.

### RNAseq on adult carcasses

Four days old mated female flies were collected in the morning and dissected in PBS. The abdomen was separated, and the gonads, the gut and the bulk of Malpighian tubes were removed, leaving the “carcass”, consisting of the abdominal fat body attached to the body wall, as well as any hemolymph, oenocytes, neurons and fragments of Malpighian tubes that remained embedded in the fat body. Precise dissection of the adult fat body without disrupting it is very difficult, which is why carcass is typically used instead (e.g. modEncode project, (Brown, et al. 2014)). From each of the 12 populations raised on either larval diet we collected one sample of 10 carcasses, i.e., 24 samples in total. The samples were snap-frozen in liquid nitrogen and stored at –80°C.

RNA was extracted from the carcass samples using “Total RNA Purification Plus” by Norgen Biotek (#48300, 48400). cDNA libraries (TrueSeq Standard RNA) were generated and sequenced on two lanes of Illumina HiSeq4000 (single read, 150 bp) by the Genomic Technologies Facility of the University of Lausanne following manufacturer’s protocols. Reads were mapped (pseudoaligned) to *D. melanogaster* reference genome BDGP6.79 using *kallisto* (Bray, et al. 2016), with 22 to 31 million mapped reads per sample. Read counts per transcript feature output from *kallisto* were converted to counts per gene using *tximport* (Soneson, et al. 2016). We filtered out genes with very low expression in that we only retained genes with read count per million greater than 2 in at least 6 samples. We further detected 45 pairs of genes with identical counts across all samples; from each pair we retained only the gene with a lower FBgn number, leaving 8701 unique genes.

Differential expression analysis was performed with *limma-voom* (Law, et al. 2014; Ritchie, et al. 2015), with evolutionary regime (Selected versus Control), with *EdgeR* normalization (Robinson, et al. 2010) implicit in the voom algorithm. Larval diet (poor versus standard) and their interaction were the fixed factors in the *limma* model; replicate population was a random factors modelled with *duplicateCorrelation* function of *limma*. Adjustment of *P*-values for multiple comparison was performed using Storey’s FDR *q*-values (Storey and Tibshirani 2003) as implemented in procedure MULTTEST option PFDR of SAS/STAT software v. 9. 4 (Copyright © 2002-2012 by SAS Institute Inc., Cary, NC, USA). Genes with expression different at *q* < 0.1 were considered as significant for enrichment analyses. GO-term enrichment analysis was carried out with bioconductor 3.12 package *topGO* v. 2.42.0 (Alexa and Rahnenfuhrer 2021), using the *weight* method and Fisher’s exact test. Because the tests for enrichment of different GO-terms are highly non-independent, no meaningful method for calculating FDR exists (Alexa and Rahnenfuhrer 2021); therefore, we report uncorrected *P*-values, focusing our interpretation on the top GO terms with *P* < 0.01.

Because the FDR threshold focuses on minimizing false positives, it leaves out many genes that have truly differ in expression. Thus, the number of genes that pass the FDR threshold greatly underestimates the number of genes that truly differ in expression, and the relationship between the two depends greatly statistical power. Therefore, we also estimated the number of genes that differed in expression due to each factor in the analysis as the total number of genes minus the estimated number of “true nulls” (Storey and Tibshirani 2003).

To study to which degree the gene expression profiles of flies from the two regimes raised on the two larval diets were separable in a multivariate space we performed principal component analysis (PCA) on the correlation matrix of the log-normalized expression data for all genes. To test for the separation of the samples in this multivariate space we performed a multivariate analysis of variance (MANOVA) on the PCA scores, with regime, diet and their interaction as the factors (using procedure GLM of SAS/STAT software).

### Larval gene expression analysis

The comparison of adult to larval gene expression differences was based on previously published data from a RNAseq study of whole larvae from the Selected and Control populations after about 190 generations of experimental evolution. All larvae were raised on the poor diet in that experiment, in either germ-free state or colonized with a single microbiota strain at the high concentration of about 10^8^ CFU (Erkosar, et al. 2017). Thus, the bacterial inoculation used for adult RNAseq (feces suspension containing about 10^3^ CFU) was intermediate between the two larval treatments. To maximize the compatibility of the adult and larval data sets we remapped the larval reads using the same *kallisto* algorithm as for adult carcasses, and analyzed the expression of the 11475 genes with the same *limma-voom* approach, with evolutionary regime, microbiota treatment and their interaction as the factors. We used the main effect of evolutionary regime from this study for the comparison with the adult results; using just data from the microbiota-colonized treatment led to qualitatively similar conclusions.

### Broad-scale targeted metabolomics

Metabolite abundance was measured using multiple pathway targeted analysis in the carcasses of 4 days old mated females obtained as described above. From each population raised on each larval diet we obtained two samples of 10 carcasses (“fed flies”). Two further samples of 10 carcasses per population and larval diet were obtained from females subject to 24 h of starvation on nutrient-deprived agarose (“starved flies”), thus resulting in a 3-way design (2 evolutionary regimes each with 6 populations × 2 current larval diets × fed vs starved flies, with 2 biological replicates, for a total of 96 samples, the number limited by cost). The number of samples was limited by the costs of metabolome analysis. The two samples for each population, larval diet and adult starvation treatment (i.e., fed or starved) were dissected by two experimenters, resulting in two experimental batches of 48 samples each. The two batches were also processed for metabolite extraction and analysis separately, on different days.

Extracted samples were analyzed by Hydrophilic Interaction Liquid Chromatography coupled to tandem mass spectrometry (HILIC - MS/MS) in both positive and negative ionization modes using a 6495 triple quadrupole system (QqQ) interfaced with 1290 UHPLC system (Agilent Technologies). Data were acquired using two complementary chromatographic separations in dynamic multiple reaction monitoring mode (dMRM) as previously described (van der Velpen, et al. 2019; Medina, et al. 2020). Data were processed using MassHunter Quantitative Analysis (for QqQ, version B.07.01/Build 7.1.524.0, Agilent Technologies). Signal intensity drift correction was performed on the pooled QC samples and metabolites with CV > 30% were discarded (Dunn, et al. 2011; Tsugawa, et al. 2014; Broadhurst, et al. 2018). In addition, a series of diluted quality controls (dQC) were used to evaluate the linearity of metabolite response; peaks with correlation to dilution factor R^2^ < 0.75 were discarded.

### Metabolome analysis

The peak area data for each compound were log-transformed, zero-centered and Pareto-scaled separately for the two experimental batches. To test for differential abundance of single compounds we fitted general mixed models (GMM) to these Pareto-scaled relative metabolite values using procedure MIXED of SAS/STAT software v. 9.4 (Copyright © 2002-2012 by SAS Institute Inc., Cary, NC, USA). For each compound we first fitted a full model, with evolutionary regime (Selected versus Control), larval diet (poor versus standard), starvation treatment (starved versus fed) and all their interactions as fixed factors, and population nested within regime, diet × population and starvation × population as random factors. Inspecting the residuals from this model we detected 13 data points (out of 10848) with externally Studentized residuals greater in magnitude than ±4.0; these data points were removed as outliers and the model was refitted.

The residuals for three of the 113 compounds (creatine, hydroxykynurenine and trehalose) deviated from normality at 10% FDR (Wilk-Shapiro test); we still report the results for these three compounds but they should be treated with caution. Because the main focus of the paper is on the metabolism of flies in their normal fed state, we also analyzed the data from the 48 samples obtained from fed flies separately, fitting GMM with regime, diet and their interaction as well as experimental batch as fixed effects, and population nested in regime and diet × population interaction as random effects. The fixed factors in the GMM were tested with type 3 *F-*tests, using Satterthwaite method to estimate the denominator degrees of freedom. *P*-values for each factor were adjusted for multiple comparison as for differentially expressed genes. To illustrate the effects of the experimental factors (in particular, the interaction between the effects of evolutionary regime and starvation; Figure 6B), for some metabolites we plotted the estimated marginal means from the GMM. Because the design was balanced and the error variance was estimated from the model, all marginal means had the same standard error; to reduce the clutter in the plots we plotted this common standard error as a single error bar rather than adding such bars to each symbol representing the mean.

To visualize and test for multivariate differentiation of the metabolome we performed PCA on log-abundances of the 113 metabolites. The values from the two batches were averaged before this analysis so there was one point per population × diet combination × starvation treatment. As for gene expression patterns, sample scores from the PCA were tested for the effects of experimental factors and their interaction in a MANOVA implemented in procedure GLM of SAS/STAT software.

We also explored the relationship between the evolutionary change in metabolite abundance and the response to starvation for several categories of compounds. We specifically examined if the difference in metabolite abundance between Selected and Control flies raised on poor food, in their normal fed state, was correlated (Pearson’s *r*) with the difference between starved and fed Control flies raised on the standard diet. These two variables are functions of non-overlapping sets of measurements, thus avoiding spurious correlations due to non-independence of errors.

### Fecundity and ovariole number

Fecundity was only assayed in females raised on poor larval diet, as described above. Male genotype and in particular the seminal fluid proteins males transfer to females during mating affect the short-term fecundity of the female. To ensure that any differences in egg number among populations are driven by female physiology and not by differences between males to which those females were mated, females from all populations were mated to males from a single laboratory population, originally collected from another site in Switzerland (Valais) in 2007. From each population, we collected 25-35 virgin females at the peak day of emergence and transferred them to standard food. Three days later we split the females in replicates of 10-12 females (2-3 replicates per population depending on the total number of females remaining alive) and placed them together with 10-12 young males on orange juice-agar medium supplemented with live yeast; they were allowed to feed and mate for 24 h. Subsequently males were discarded (to prevent potential fecundity reduction due to male harassment) and the groups of females allowed to oviposit overnight in new bottles with fresh orange juice-agar supplemented with live yeast. Eggs were washed from the medium surface with tap water, collected on a fine nylon mesh and transferred to a well of a 12-well cell culture plate containing 3 mL of 1% sodium dodecyl sulfate (SDS, Sigma) to facilitate egg dispersion. A photograph of each well was taken under a Leica stereomicroscope with a Canon 60D camera with manual exposure programming, automatic white balance, 3.2 sec of exposure time and ISO 400.

The number of eggs was estimated automatically with Codicount ImageJ plugin (https://www6.paca.inrae.fr/institut-sophia-agrobiotech_eng/CODICOUNT), following Perez (2017). This approach is based on automatically quantifying the area corresponding to the eggs on the image, based on color contrast between the eggs and the background. This total egg area was then converted to an estimate of the number of eggs by dividing it by the area of a single egg, estimated from a separate sample (the same standard egg area was used for all images). Thus estimated number of eggs was divided by the number of females in a particular replicate. In one replicate one of ten females was found dead at the end of oviposition; for the analysis we set the number of females in this bottle at 9.5.

Flies were raised on the poor larval diet. While the number of ovarioles is fixed by the time of emergence, they are better visible and easier to count if filled with developing eggs (Bergland, et al. 2008). We therefore allowed 25 freshly emerged females to mate with 25 males on standard diet for three days, and subsequently to feed on ad libitum live yeast for two days, before collecting and storing them at –80°C until dissections. Six females from each population were haphazardly chosen for ovary dissection. We dissected the ovaries by separating the abdomen and cutting its posterior end, opening the abdomen laterally and removing the ovaries. We then dipped the ovaries briefly in a solution of crystal violet to improve visual contrast, opened them and counted the number of ovarioles in each ovary. Dissections and counting were done blindly with respect to the identity of the sample. To normalize the fecundity of each population relative to ovariole number we divided the mean number of eggs per female in this population by the mean number of ovarioles.

The number of replicates was based on practical considerations, no formal power analysis has been conducted. The estimation of egg number of counting of ovarioles was performed blindly with respect to the identity of the sample. Egg number and ovariole number per female were analyzed with a GMM, with regime as a fixed factor and population nested in regime as a random factor. For the number of eggs per ovariole we only had one data point per population, these were compared between regimes with a one-way linear model.

## Acknowledgements

This project has been supported by the Swiss National Science Foundation (grant 31003A_162732 to T.J.K.) and by research funding from the University of Lausanne. We thank Tony Teav at Metabolomics Platform at UNIL for his contribution to sample preparation and data processing, and three reviewers for their constructive comments on a previous version of the manuscript.

## Author contributions

B.E. and T.J.K designed the study; C.D. F.C., H.G.A. and J.I. B.E. contributed to design of experiments. B.E., C.D., F.C. L.S. and L.K. carried out the experiments. H.G.A. and J.I. performed metabolite analysis. B.E. and T.J.K. analyzed the data and wrote the paper, with contributions from other authors.

### Competing interests

The authors declare no competing interests.

## Data availability

The raw and processed data from the RNAseq on adult carcasses are available from NCBI GEO (accession number GSE193105). Raw data for the previously published larval RNAseq are available from NCBI SRA (accession numbers SAMN07723150-SAMN07723173). Previously published larval metabolome data are available as supplementary material to Cavigliasso, et al. (2023). The adult metabolite abundance data are provided in Supplementary file 8), fecundity and ovariole data in Figure 7—source data 1.

---

**Figure 1 – figure supplement 1.**
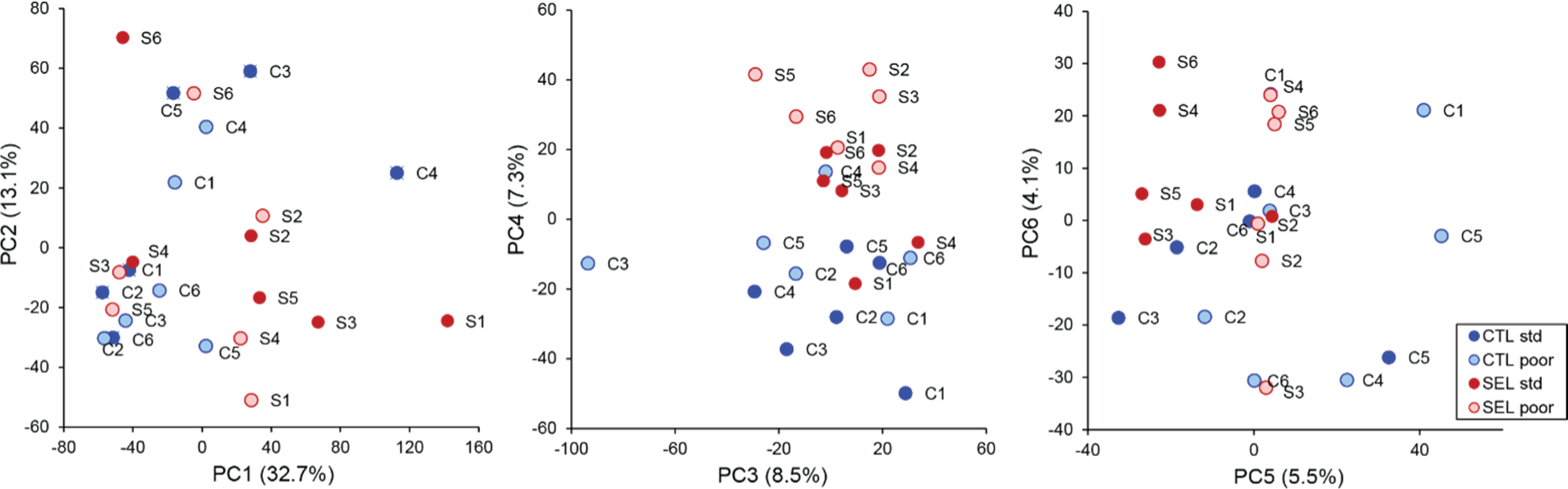
Plots of the first six principal components of the gene expression levels of adults from the Selected and Control populations raised on the standard and poor diet (Figure 1—source data 1).

**Figure 5 – figure supplement 1.**
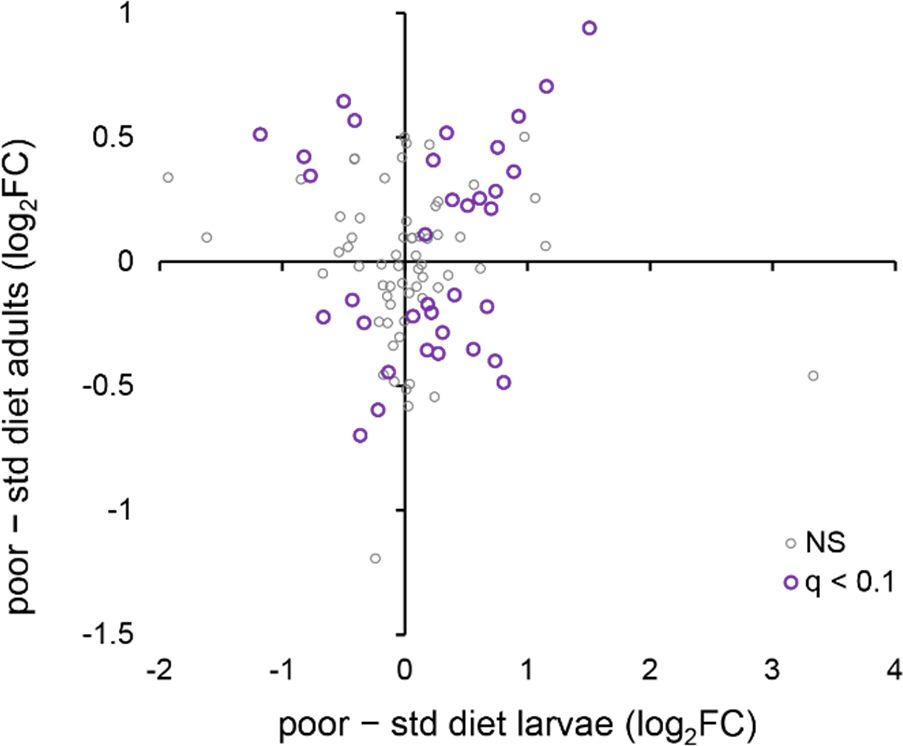
The relationship between the phenotypically plastic responses of the metabolomes of larvae and adults to larval diet. The plotted values are main effect estimates from the statistical models (i.e., they are averaged across the Selected and Control populations). *q* < 0.1 refers to metabolites that pass the 10% FDR threshold for both stages; NS are the remaining metabolites (Figure 5—source data 1, sheet C).

**Figure 6—figure supplement 1.**
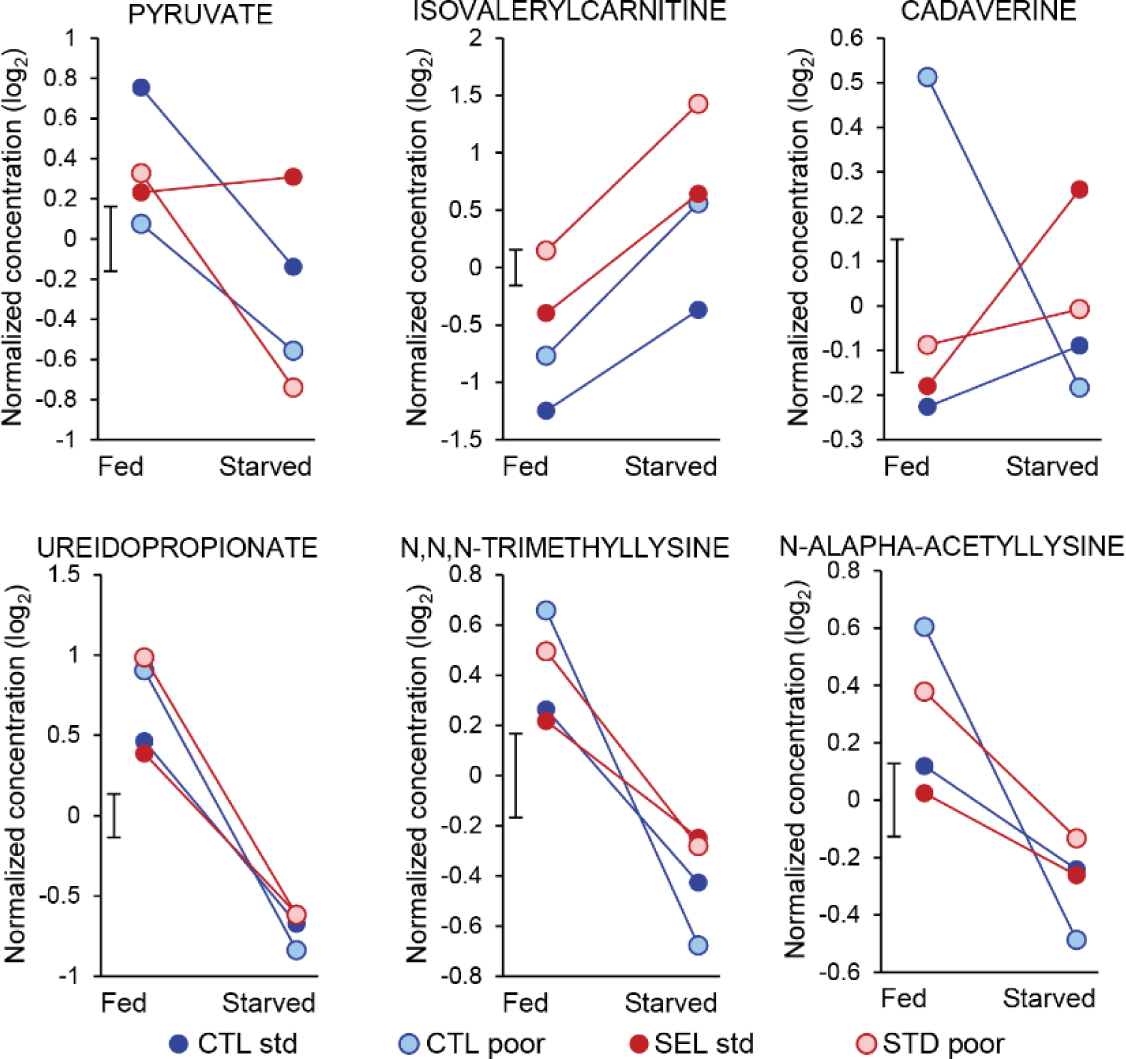
Metabolites showing a significant 3-way interaction evolutionary regime × larval diet × starvation (pyruvate) or the 2-way interaction between larval diet and starvation treatment (the remaining five compounds). *q* < 0.1 was used as the significance threshold. Symbols indicate least-square means obtained from the full factorial general mixed model (Figure 6—source data 2). The bar next to the Y-axis indicates the standard error of the means (identical for all treatments as they were estimated from the model).

## List of supplementary files

Figure 1—source data 1: PCA analysis of adult gene expression: the first six PC scores of all populations on each diet.

Figure 2—source data 1: PCA analysis of adult metabolite abundance in fed flies: the first four PC scores of all populations

Figure 4—source data 1: The estimated main effects of diet (the plastic response) and of evolutionary regime (the evolutionary change) on adult gene expression and metabolite abundance

Figure 5—source data 1: The relationship between evolutionary change in adults and the corresponding change in the larvae

Figure 6—source data 1: PCA analysis of metabolite abundance in fed and starved flies: the first four PC scores of all populations under all conditions

Figure 6—source data 2: Abundance patters of metabolites with interactions

Figure 6—source data 3: Relationship between response to starvation and the evolutionary change Figure 7—source data 1: Data on fecundity and ovariole number

Supplementary file 1: Results of univariate analysis of gene expression in adult female carcass performed in limma-voom.

Supplementary file 2: Results of GO-term enrichment for “biological process” on genes expressed in female adult carcass. (A) Genes differentially expressed between Selected and Control flies. (B) Genes differentially expressed in flies raised on standard versus poor larval diet. (C) Genes differentially expressed between Selected and Control populations both in adults and larvae. (D) Genes differentially expressed between Selected and Control populations in adults but not in larvae.

Supplementary file 3: Results of univariate analysis of metabolome with general mixed model. (A) Least-square means for all combinations of the three experimental factors: evolutionary regime (Control versus Selected), larval diet (standard versus poor) and adult starvation treatment (fed versus starved). The LS means and standard errors (SE) were obtained from the full GMM model. They are expressed on a zero-centered log2 scale. (B) Least-square means contrasts corresponding to Figure 2B: Regime(Fed) = Selected – Control in fed condition; Diet(Fed) = flies in fed condition raised on standard diet – flies in fed condition raised on poor diet; Starv = starved flies – fed flies across both regimes and diets. (C) Tests of significance (df, F, nominal P, q = adjusted P) from separate analysis on flies in fed condition only. (D) Tests of significance from the full model on both fed and starved flies. Denominator degrees of freedom (df) were estimated with the Satterthwaite method and thus vary across compounds depending on the magnitude of variance components corresponding to random factors.

Supplementary file 4: Results of Joint Pathway Analysis on metabolites significantly different between Selected and Control flies (in fed state).

Supplementary file 5: Genes that show differential expression between Selected and Control populations (at raw *P* < 0.05) at both larval and adult stage.

Supplementary file 6: Analysis of the effect of evolutionary regime on the expression of genes involved in amino acid metabolism and purine synthesis in the larvae.

Supplementary file 7: MANOVA on the first two principal component scores from the PCA on metabolite abundance in fed and starved flies.

Supplementary file 8: Original data on metabolite abundance. The table includes the original estimates of peak area (in arbitrary units) normalized to the sample protein content, and log10-transformed, zero-centered and Pareto-scaled values used for the analysis (zero-centering and Pareto-scaling was done separately for each batch). The table includes the 13 outlier data points that were removed from the final analysis because their Studentized residuals exceeded ±4, these outliers are identified in the last column.

## Notes

https://www.ncbi.nlm.nih.gov/geo/query/acc.cgi?acc=GSE193105

https://www.ncbi.nlm.nih.gov/bioproject/PRJNA412704

https://academic.oup.com/evlett/article/7/4/273/7170067#supplementary-data

### Competing Interest Statement

The authors have declared no competing interest.

### Summary of Updates

This is a largely rewritten ms with a different structure of Results. The Results contain several new analyses and in particular an analysis of the relationship between differences due to evolutionary regime at the larval and the adult stage.

## References

Agnoux AM, Antignac JP, Simard G, Poupeau G, Darmaun G, Parnet P, Alexandre-Gouabau MC. 2014. Time window-dependent effect of perinatal maternal protein restriction on insulin sensitivity and energy substrate oxidation in adult male offspring. American Journal of Physiology-Regulatory, Integrative and Comparative Physiology 307:R184–R197.

Akitake Y, Katsuragi S, Hosokawa M, Mishima K, Ikeda T, Miyazato M, Hosoda H. 2015. Moderate maternal food restriction in mice impairs physical growth, behavior, and neurodevelopment of offspring. Nutrition Research 35:76–87.

Alexa A, Rahnenfuhrer J. 2021. topGO: Enrichment Analysis for Gene Ontology. R package version 2.46.0.

Antikainen H, Driscoll M, Haspel G, Dobrowolski R. 2017. TOR-mediated regulation of metabolism in aging. Aging Cell 16:1219–1233.

Baldwin JM. 1896. A new factor in evolution. The American Naturalist 30:441–451,536-553.

Bateson P, Barker D, Clutton-Brock T, Deb D, D’Udine B, Foley RA, Gluckman P, Godfrey K, Kirkwood T, Lahr MM, et al. 2004. Developmental plasticity and human health. Nature 430:419–421.

Bergland AO, Genissel A, Nuzhdin SV, Tatar M. 2008. Quantitative trait loci affecting phenotypic plasticity and the allometric relationship of ovariole number and thorax length in Drosophila melanogaster. Genetics 180:567–582.

Bray NL, Pimentel H, Melsted P, Pachter L. 2016. Near-optimal probabilistic RNA-seq quantification. Nature Biotechnology 34:525–527.

Broadhurst D, Goodacre R, Reinke SN, Kuligowski J, Wilson ID, Lewis MR, Dunn WB. 2018. Guidelines and considerations for the use of system suitability and quality control samples in mass spectrometry assays applied in untargeted clinical metabolomic studies. Metabolomics 14:72.

Brown JB, Boley N, Eisman R, May GE, Stoiber MH, Duff MO, Booth BW, Wen J, Park S, Suzuki AM, et al. 2014. Diversity and dynamics of the Drosophila transcriptome. Nature 512:393–399.

Carlsson H, Ivimey-Cook E, Duxbury EML, Edden N, Sales K, Maklakov AA. 2021. Ageing as “early-life inertia”: Disentangling life-history trade-offs along a lifetime of an individual. Evolution Letters 5:551–564.

Cavigliasso F, Dupuis C, Savary L, Spangenberg JE, Kawecki TJ. 2020. Experimental evolution of post-ingestive nutritional compensation in response to a nutrient-poor diet. Proceedings of the Royal Society B: Biological Sciences 287:20202684.

Cavigliasso F, Savary L, Spangenberg JE, Gallart-Ayala H, Ivanisevic J, Kawecki TJ. 2023. Experimental evolution of metabolism under nutrient restriction: enhanced amino acid catabolism and a key role of branched-chain amino acids. Evolution Letters 7:273–284.

Cohen E, Sawyer JK, Peterson NG, Dow JAT, Fox DT. 2020. Physiology, Development, and Disease Modeling in the Drosophila Excretory System. Genetics 214:235–264.

Collet J, Fellous S. 2019. Do traits separated by metamorphosis evolve independently? Concepts and methods. Proceedings of the Royal Society B-Biological Sciences 286.

Davis K, Chamseddine D, Harper JM. 2016. Nutritional limitation in early postnatal life and its effect on aging and longevity in rodents. Experimental Gerontology 86:84–89.

de Brito Alves JL, Nogueira VO, de Oliveira GB, da Silva GSF, Wanderley AG, Leandro CG, Costa-Silva JH. 2014. Short- and long-term effects of a maternal low-protein diet on ventilation, O2/CO2 chemoreception and arterial blood pressure in male rat offspring. British Journal of Nutrition 111:606–615.

de Magalhaes JP. 2012. Programmatic features of aging originating in development: aging mechanisms beyond molecular damage? Faseb Journal 26:4821–4826.

de Rooij SR, Wouters H, Yonker JE, Painter RC, Roseboom TJ. 2010. Prenatal undernutrition and cognitive function in late adulthood. Proceedings of the National Academy of Sciences 107:16881–16886.

Dunn WB, Broadhurst D, Begley P, Zelena E, Francis-McIntyre S, Anderson N, Brown M, Knowles JD, Halsall A, Haselden JN, et al. 2011. Procedures for large-scale metabolic profiling of serum and plasma using gas chromatography and liquid chromatography coupled to mass spectrometry. Nature Protocols 6:1060–1083.

Erkosar B, Kolly S, van der Meer JR, Kawecki TJ. 2017. Adaptation to chronic nutritional stress leads to reduced dependence on microbiota in *Drosophila*. mBio 8:e01496–01417.

Gems D, Partridge L. 2013. Genetics of Longevity in Model Organisms: Debates and Paradigm Shifts. In: Julius D, editor. Annual Review of Physiology, Vol 75. p. 621–644.

Ghalambor CK, Hoke KL, Ruell EW, Fischer EK, Reznick DN, Hughes KA. 2015. Non-adaptive plasticity potentiates rapid adaptive evolution of gene expression in nature. Nature 525:372–375.

Grandison RC, Piper MDW, Partridge L. 2009. Amino-acid imbalance explains extension of lifespan by dietary restriction in Drosophila. Nature 462:1061–U1121.

Gu X, Jouandin P, Lalgudi PV, Binari R, Valenstein ML, Reid MA, Allen AE, Kamitaki N, Locasale JW, Perrimon N, et al. 2022. Sestrin mediates detection of and adaptation to low-leucine diets in Drosophila. Nature 608:209–216.

Gupta V, Frank AM, Matolka N, Lazzaro BP. 2022. Inherent constraints on a polyfunctional tissue lead to a reproduction-immunity tradeoff. Bmc Biology 20:127.

Hamilton WD. 1966. The moulding of senescence by natural selection. Journal of Theoretical Biology 12:12–45.

Hollis B, Houle D, Yan Z, Kawecki TJ, Keller L. 2014. Evolution under monogamy feminizes gene expression in Drosophila melanogaster. Nature Communications 5:3482.

Huang Y, Agrawal AF. 2016. Experimental Evolution of Gene Expression and Plasticity in Alternative Selective Regimes. Plos Genetics 12:e1006336.

Josephs EB, Van Etten ML, Harkess A, Platts A, Baucom RS. 2021. Adaptive and maladaptive expression plasticity underlying herbicide resistance in an agricultural weed. Evolution Letters 5:432–440.

Kanehisa M, Goto S. 2000. KEGG: Kyoto Encyclopedia of Genes and Genomes. Nucleic Acids Research 28:27–30.

Kapila R, Kashyap M, Gulati A, Narasimhan A, Poddar S, Mukhopadhaya A, Prasad NG. 2021. Evolution of sex-specific heat stress tolerance and larval Hsp70 expression in populations of Drosophila melanogaster adapted to larval crowding. Journal of Evolutionary Biology 34:1376–1385.

Kastanos EK, Woldman YY, Appling DR. 1997. Role of mitochondrial and cytoplasmic serine hydroxymethyltransferase isozymes in de novo purine synthesis in Saccharomyces cerevisiae. Biochemistry 36:14956–14964.

Kawecki TJ, Erkosar B, Dupuis C, Hollis B, Stillwell RC, Kapun M. 2021. The genomic architecture of adaptation to larval malnutrition points to a trade-off with adult starvation resistance in *Drosophila*. Molecular Biology and Evolution 38:2732–2749.

Klepsatel P, Knoblochova D, Girish TN, Dircksen H, Galikova M. 2020. The influence of developmental diet on reproduction and metabolism inDrosophila. Bmc Evolutionary Biology 20.

Kolss M, Vijendravarma RK, Schwaller G, Kawecki TJ. 2009. Life history consequences of adaptation to larval nutritional stress in *Drosophila*. Evolution 63:2389–2401.

Laland KN, Uller T, Feldman MW, Sterelny K, Müller GB, Moczek A, Jablonka E, Odling-Smee J. 2015. The extended evolutionary synthesis: its structure, assumptions and predictions. Proceedings of the Royal Society B: Biological Sciences 282:20151019.

Lande R, Arnold SJ. 1983. The measurement of selection on correlated characters. Evolution 37:1210–1226.

Law CW, Chen Y, Shi W, Smyth GK. 2014. voom: precision weights unlock linear model analysis tools for RNA-seq read counts. Genome Biology 15:R29.

Li S, Yu X, Feng Q. 2019. Fat Body Biology in the Last Decade. Annual Review of Entomology 64:315–333.

Lillycrop KA, Burdge GC. 2015. Maternal diet as a modifier of offspring epigenetics. Journal of Developmental Origins of Health and Disease 6:88–95.

Lukaszewski MA, Eberle D, Vieau D, Breton C. 2013. Nutritional manipulations in the perinatal period program adipose tissue in offspring. American Journal of Physiology-Endocrinology and Metabolism 305:E1195–E1207.

May CM, van den Heuvel J, Doroszuk A, Hoedjes KM, Flatt T, Zwaan BJ. 2019. Adaptation to developmental diet influences the response to selection on age at reproduction in the fruit fly. Journal of Evolutionary Biology 32:425–437.

May CM, Zwaan BJ. 2017. Relating past and present diet to phenotypic and transcriptomic variation in the fruit fly. Bmc Genomics 18.

Medawar PB. 1952. An unsolved problem of biology. London: H. K. Lewis.

Medina J, van der Velpen V, Teav T, Guitton Y, Gallart-Ayala H, Ivanisevic J. 2020. Single-Step Extraction Coupled with Targeted HILIC-MS/MS Approach for Comprehensive Analysis of Human Plasma Lipidome and Polar Metabolome. Metabolites 10:495.

Moczek AP, Sultan S, Foster S, Ledón-Rettig C, Dworkin I, Nijhout HF, Abouheif E, Pfennig DW. 2011. The role of developmental plasticity in evolutionary innovation. Proceedings of the Royal Society B: Biological Sciences 278:2705–2713.

Moran NA. 1994. Adaptation and constraint in the complex life cycles of animals. Annual Review of Ecology and Systematics 25:573–600.

Neel JV. 1962. Diabetes mellitus: a “thrifty” genotype rendered detrimental by “progress”? American Journal of Human Genetics 14:353–362.

O’Brien DM, Min K-J, Larsen T, Tatar M. 2008. Use of stable isotopes to examine how dietary restriction extends Drosophila lifespan. Current Biology 18:R155–R156.

Partridge L, Barton NH. 1993. Optimality, mutation and the evolution of ageing. Nature 362:305–311.

Perez G. 2017. Une méthode d’analyse d’image automatique pour quantifier rapidement les nombres d’œufs et les taux de parasitisme chez Trichogramma sp. In. Cahiers techniques de l’INRA. Innovations entomologiques: du laboratoire au champ: INRA. p. 135–142.

Pigliucci M, Murren CJ. 2003. Genetic assimilation and a possible evolutionary paradox: can macroevolution sometimes be so fast as to pass us by? Evolution 57:1455–1464.

Piper MDW, Soultoukis GA, Blanc E, Mesaros A, Herbert SL, Juricic P, He X, Atanassov I, Salmonowicz H, Yang M, et al. 2017. Matching Dietary Amino Acid Balance to the In Silico-Translated Exome Optimizes Growth and Reproduction without Cost to Lifespan. Cell Metabolism 25:610–621.

Pischedda A, Chippindale AK. 2006. Intralocus sexual conflict diminishes the benefits of sexual selection. Plos Biology 4:2099–2103.

Prentice AM, Rayco-Solon P, Moore SE. 2005. Insights from the developing world: thrifty genotypes and thrifty phenotypes. Proceedings of the Nutrition Society 64:153–161.

Rice WR. 1984. Sex chromosomes and the evolution of sexual dimorphism. Evolution 38:735–742.

Ritchie ME, Phipson B, Wu D, Hu Y, Law CW, Shi W, Smyth GK. 2015. limma powers differential expression analyses for RNA-sequencing and microarray studies. Nucleic Acids Research 43:e47–e47.

Robinson MD, McCarthy DJ, Smyth GK. (Robinson2010 co-authors). 2010. edgeR: a Bioconductor package for differential expression analysis of digital gene expression data. Bioinformatics 26.

Rolff J, Johnston PR, Reynolds S. 2019. Complete metamorphosis of insects. Philosophical Transactions of the Royal Society B: Biological Sciences 374:20190063.

Roseboom T, de Rooij S, Painter R. 2006. The Dutch famine and its long-term consequences for adult health. Early Hum Dev 82:485–491.

Salway JG. 2018. The Krebs Uric Acid Cycle: A Forgotten Krebs Cycle. Trends in Biochemical Sciences 43:847–849.

Schaerli Y, Jiménez A, Duarte JM, Mihajlovic L, Renggli J, Isalan M, Sharpe J, Wagner A. 2018. Synthetic circuits reveal how mechanisms of gene regulatory networks constrain evolution. Molecular Systems Biology 14:e8102.

Scheiner SM. 1993. Genetics and evolution of phenotypic plasticity. Annual Review of Ecology and Systematics 24:35–68.

Schmidt PS, Matzkin L, Ippolito M, Eanes WF. 2005. Geographic variation in diapause incidence, life-history traits, and climatic adaptation in *Drosophila melanogaster*. Evolution 59:1721–1732.

Scott RC, Schuldiner O, Neufeld TP. 2004. Role and Regulation of Starvation-Induced Autophagy in the Drosophila Fat Body. Developmental Cell 7:167–178.

Shenoi VN, Ali SZ, Prasad NG. 2016. Evolution of increased adult longevity in Drosophila melanogaster populations selected for adaptation to larval crowding. Journal of Evolutionary Biology 29:407–417.

Soneson C, Love M, Robinson M. 2016. Differential analyses for RNA-seq: transcript-level estimates improve gene-level inferences [version 2; peer review: 2 approved]. F1000Research 4.

Sorrells TR, Booth LN, Tuch BB, Johnson AD. 2015. Intersecting transcription networks constrain gene regulatory evolution. Nature 523:361–365.

Stanley-Samuelson DW, Jurenka RA, Cripps C, Blomquist GJ, Derenobales M. 1988. Fatty acids in insects - composition, metabolism and biological significance. Archives of Insect Biochemistry and Physiology 9:1–33.

Storey JD, Tibshirani R. 2003. Statistical significance for genomewide studies. Proceedings of the National Academy of Sciences of the United States of America 100:9440–9445.

Szostaczuk N, van Schothorst EM, Sanchez J, Priego T, Palou M, Bekkenkamp-Grovenstein M, Faustmann G, Obermayer-Pietsch B, Tiran B, Roob JM, et al. 2020. Identification of blood cell transcriptome-based biomarkers in adulthood predictive of increased risk to develop metabolic disorders using early life intervention rat models. Faseb Journal 34:9003–9017.

Tsugawa H, Kanazawa M, Ogiwara A, Arita M. 2014. MRMPROBS suite for metabolomics using large-scale MRM assays. Bioinformatics 30:2379–2380.

van der Velpen V, Teav T, Gallart-Ayala H, Mehl F, Konz I, Clark C, Oikonomidi A, Peyratout G, Henry H, Delorenzi M, et al. 2019. Systemic and central nervous system metabolic alterations in Alzheimer’s disease. Alzheimer’s research & therapy. doi: 10.1186/s13195-019-0551-7

Veltsos P, Fang YX, Cossins AR, Snook RR, Ritchie MG. 2017. Mating system manipulation and the evolution of sex-biased gene expression in Drosophila. Nature Communications 8.

Vijendravarma RK, Kawecki TJ. 2013. Epistasis and maternal effects in experimental adaptation to chronic nutritional stress in *Drosophila*. Journal of Evolutionary Biology 26:2566–2580.

Vijendravarma RK, Narasimha S, Chakrabarti S, Babin A, Kolly S, Lemaitre B, Kawecki TJ. 2015. Gut physiology mediates a trade-off between adaptation to malnutrition and susceptibility to food-borne pathogens. Ecology Letters 18:1078–1086.

Vijendravarma RK, Narasimha S, Kawecki TJ. 2012. Chronic malnutrition favours smaller critical size for metamorphosis initiation in *Drosophila melanogaster*. Journal of Evolutionary Biology 25:288–292.

Vijendravarma RK, Narasimha S, Kawecki TJ. 2010. Effects of parental larval diet on egg size and offspring traits in *Drosophila*. Biology Letters 6:238–241.

Wagner A. 2011. Genotype networks shed light on evolutionary constraints. Trends in Ecology & Evolution 26:577–584.

Wolfson RL, Chantranupong L, Saxton RA, Shen K, Scaria SM, Cantor JR, Sabatini DM. 2016. Sestrin2 is a leucine sensor for the mTORC1 pathway. Science 351:43–48.

Xia J, Psychogios N, Young N, Wishart DS. 2009. MetaboAnalyst: a web server for metabolomic data analysis and interpretation. Nucleic Acids Research 37:W652–W660.

Yampolsky LY, Glazko GV, Fry JD. 2012. Evolution of gene expression and expression plasticity in long-term experimental populations of Drosophila melanogaster maintained under constant and variable ethanol stress. Molecular Ecology 21:4287–4299.

